# Hepatocellular carcinoma-associated *AXIN1* mutations drive low levels of Wnt/β-catenin pathway activity that allow for niche-independent growth and YAP/TAZ signaling

**DOI:** 10.1101/2024.11.28.625941

**Authors:** Anton J. Venhuizen, Yvanka van Os, Milo L. Kaptein, Marleen T. Aarts, Despina Xanthakis, Ingrid Jordens, Madelon M. Maurice

**Affiliations:** Oncode Institute and Center for Molecular Medicine, University Medical Center Utrecht, 3584 CX, Utrecht, The Netherlands

**Keywords:** AXIN1, CTNNB1, Wnt/β-catenin signaling, YAP/TAZ signaling, hepatocellular carcinoma

## Abstract

In healthy cells, AXIN1 organizes assembly of a large destruction complex that mediates proteolysis of the transcriptional co-activator β-catenin to prevent inappropriate Wnt/β-catenin pathway activation. In hepatocellular carcinoma (HCC), *AXIN1* mutations (11%) associate with a poor-prognosis subtype that is molecularly distinct from β-catenin-mutant HCC (28-40%). How *AXIN1* deficiency drives HCC formation has remained highly debated. Here, we address this issue by introducing HCC-associated *AXIN1* and *CTNNB1* mutations in human liver cancer cells and liver-derived organoids. We show that different mutant *AXIN1* classes activate varying degrees of Wnt signaling, although at lower overall levels than *CTNNB1* mutations. Strikingly, premature stop codons in 5’ coding regions do not classify as knock-out mutations but drive alternative translation of an N-terminally truncated AXIN1 variant with partially retained suppressor activity. All *AXIN1* variants endow liver progenitor organoids with the capacity to grow in the absence of R-spondin and Wnt, indicative of aberrant Wnt/β-catenin pathway activation. Additionally, induced Wnt/β-catenin pathway activation inversely correlates with YAP/TAZ-mediated signaling, thus leaving higher residual YAP/TAZ activity in *AXIN1*-mutant versus *CTNNB1*-mutant cells. We conclude that *AXIN1* mutations drive physiologically relevant Wnt/β-catenin signaling in HCC, while providing a permissive environment for YAP/YAZ signaling, thereby distinguishing *AXIN1* mutations from those in *CTNNB1*.

## Introduction

Hepatocellular carcinoma (HCC) is the most prevalent form of liver cancer and the third deadliest malignancy worldwide, accounting for over 800.000 deaths per year^1^. Although HCC is a heterogeneous disease, mutational alterations that affect core components of the Wnt/β-catenin pathway are identified in a prominent fraction of patients^2^. In the healthy liver, the evolutionary conserved Wnt/β-catenin pathway governs development, homeostasis, injury-induced regeneration, and zonation^3^. Furthermore, levels of Wnt/β-catenin signaling determine metabolic identity and correlate with proliferation rates of hepatocytes that constitute the majority of liver parenchyma cells^4^. The primary outcome of Wnt/β-catenin pathway activation is stabilization of the transcriptional co-factor β-catenin, which promotes its nuclear entry and complex formation with TCF/LEF to mediate the transcription of Wnt target genes^5^. In the absence of Wnts, cytosolic β-catenin levels are constitutively targeted for proteolysis by a multiprotein β-catenin destruction complex. AXIN1, the principal coordinator of this complex, brings together the scaffold APC and kinases GSK3 and CK1 to orchestrate efficient phosphorylation, ubiquitination and proteasomal degradation of β-catenin^6^. Upon binding of Wnts to their receptors at the cell surface, the destruction complex is inactivated, resulting in β-catenin accumulation and the activation of gene programs linked to stem cell maintenance and proliferation^5^.

For HCC, the most frequent alterations in Wnt pathway components comprise activating mutations in β-catenin (gene name: *CTNNB1*, mutated in 28-40% of cases) and inactivating mutations in *AXIN1* (mutated in 11% of cases)^7^. Historically, *AXIN1* and *CTNNB1* mutations in HCC were considered functionally equivalent, as cell lines harboring mutations in either gene displayed increased complex formation between β-catenin and TCF/LEF, indicative of Wnt pathway activation^8^. Recent studies however linked *CTNNB1* and *AXIN1* mutations to molecularly distinct HCC subtypes, each with divergent pathological implications. *CTNNB1* mutations generally result in a Wnt-high expression profile, and are predominantly found in the class of ‘non-proliferative’ tumors that associate with better prognosis^9^. By contrast, *AXIN1* mutations are linked to the ‘proliferative’ Wnt-low class of more aggressive HCC tumors, hallmarked by poor differentiation. To what extent *AXIN1* mutations activate Wnt signaling in HCC, remains a highly debated issue. AXIN1-deficient mice were reported to develop liver tumors without indications of Wnt pathway activation^10^. Instead, these tumors displayed elevated levels of YAP/YAZ signaling, NOTCH pathway activation and fetal gene expression. In line with these findings, a recent study showed that YAP/TAZ-mediated signaling is indispensable for liver tumor formation in AXIN1- deficient mice^11^. Moreover, overexpression of YAP promotes expansion of progenitor-like murine hepatocytes, further linking YAP/TAZ signaling to the ‘proliferative’, dedifferentiated HCC subtype associated with *AXIN1* mutations^12^.

In contrast to these reports, we and others established a connection between *AXIN1* mutations and elevated Wnt signaling. For instance, HCC tumors harboring *AXIN1* mutations displayed upregulation of a subset of Wnt target genes^13,14^. Furthermore, several cancer-associated *AXIN1* missense mutations in the N-terminal RGS domain of AXIN1 were found to induce Wnt signaling and drive hyperplastic growth in *Drosophila* wing discs^15^. Recent studies further showed that various *AXIN1* missense and truncating mutations promote Wnt/β-catenin pathway activation and, moreover, signaling levels directly correlated with tumorigenic potential in the liver^16,17^.

In summary, *AXIN1*-mutant liver tumors depend on both β-catenin and YAP/TAZ signaling, however the relative contributions of these pathways remain disputed. While *AXIN1*-mutant HCC cell lines are generally considered Wnt-low compared to *CTNNB1*-mutant cells, a comprehensive comparison between both mutation types in HCC cells with a similar genetic background is lacking^18^. Furthermore, it is unclear whether *AXIN1* mutations endow niche factor-independent growth in the liver, as observed for other Wnt pathway mutations in various tissues, such as the colon^19^.

To address these issues, we introduced a variety of HCC-associated *AXIN1* mutations as well as activating *CTNNB1* mutations in the liver cancer cell line Huh7, HEK293T cells and mouse liver progenitor organoids. We observe that missense and truncating *AXIN1* mutations activate Wnt/β-catenin signaling, although to varying degrees and less pronounced than *CTNNB1* mutations. Unexpectedly, we found that the insertion of premature stop codons in the 5’ exonic regions of *AXIN1* do not result in knock-out but rather promote alternative translation of N- terminally truncated AXIN1 variants that display partially retained suppressor activity. Mouse liver progenitor organoids harboring *Axin1* alterations could be propagated in the absence of Wnt and R-spondin (RSPO) ligands, indicating that *Axin1* mutations enable niche-independent Wnt pathway activation in the liver. Notably, we uncover an inverse correlation between the degree of mutation-induced Wnt pathway activation and the capacity of cells to activate YAP/TAZ signaling. Our findings thus provide an explanation for why Wnt-high, *CTNNB1*-mutant tumors generally lack YAP/TAZ activity while Wnt-low, *AXIN1*-mutant tumors associate with enhanced YAP/TAZ- mediated gene transcription and a more clinically aggressive phenotype.

## Results

### Missense mutations within the RGS domain of AXIN1 drive Wnt/β-catenin signaling

To assess the roles of *CTNNB1* and *AXIN1* mutations in hepatocellular carcinoma (HCC), we examined mutational patterns in these genes as listed in the cBioPortal database (Fig. 1a-b)^20–25^. For *CTNNB1*, the majority of missense mutations (88%) locate to the N-terminal degron motif of β-catenin and thus interfere with its phosphorylation, ubiquitination and proteasomal degradation^26^. By contrast, genetic alterations in *AXIN1* are primarily truncating and missense mutations that are scattered throughout the gene. We previously demonstrated that cancer-associated single amino acid substitutions within the RGS domain of AXIN1 promote unfolding and the formation of soluble nano-aggregates, leading to impaired AXIN1 polymerization, loss of APC binding and reduced destruction complex activity^15^. We set out to assess the functional consequences of all *AXIN1* missense variants reported in the TCGA HCC database, using a β- catenin-dependent luciferase reporter assay (TOPFlash) in HEK293T cells^27^. Most AXIN1 variants carrying a mutant RGS domain induced ligand-independent activation of the Wnt/β-catenin pathway upon overexpression (Fig. 1c). Concomitantly, these mutations disrupted AXIN1 polymerization and stability, as shown by loss of cytosolic puncta formation (Fig. S1a) and diminished overall protein expression levels (Fig. 1d), in line with the previously reported L106R variant^15^. AXIN1 A120D formed the only exception, as for this variant β-catenin-mediated reporter activity was induced while puncta formation and protein levels remained unaltered (Fig 1d, S1a), in line with its decreased propensity for aggregation in comparison with the other RGS missense variants (Fig. S1b)^15^. The AXIN1 A120D mutation likely interferes with APC binding directly, given its position in the APC interaction interface (Fig. S1c)^17^. Notably, introduction of missense mutations in the AXIN1 DIX domain did not lead to ligand-independent activation of Wnt/β-catenin signaling in this model system, even though polymerization was perturbed (Fig. 1c; S1a and d). In a previous study, we showed that such variants display impaired suppressor activity when expressed at lower, endogenous concentrations^28^, indicating that these more subtle defects may be overcome by increased AXIN1 concentrations due to overexpression.

**Figure 1.**
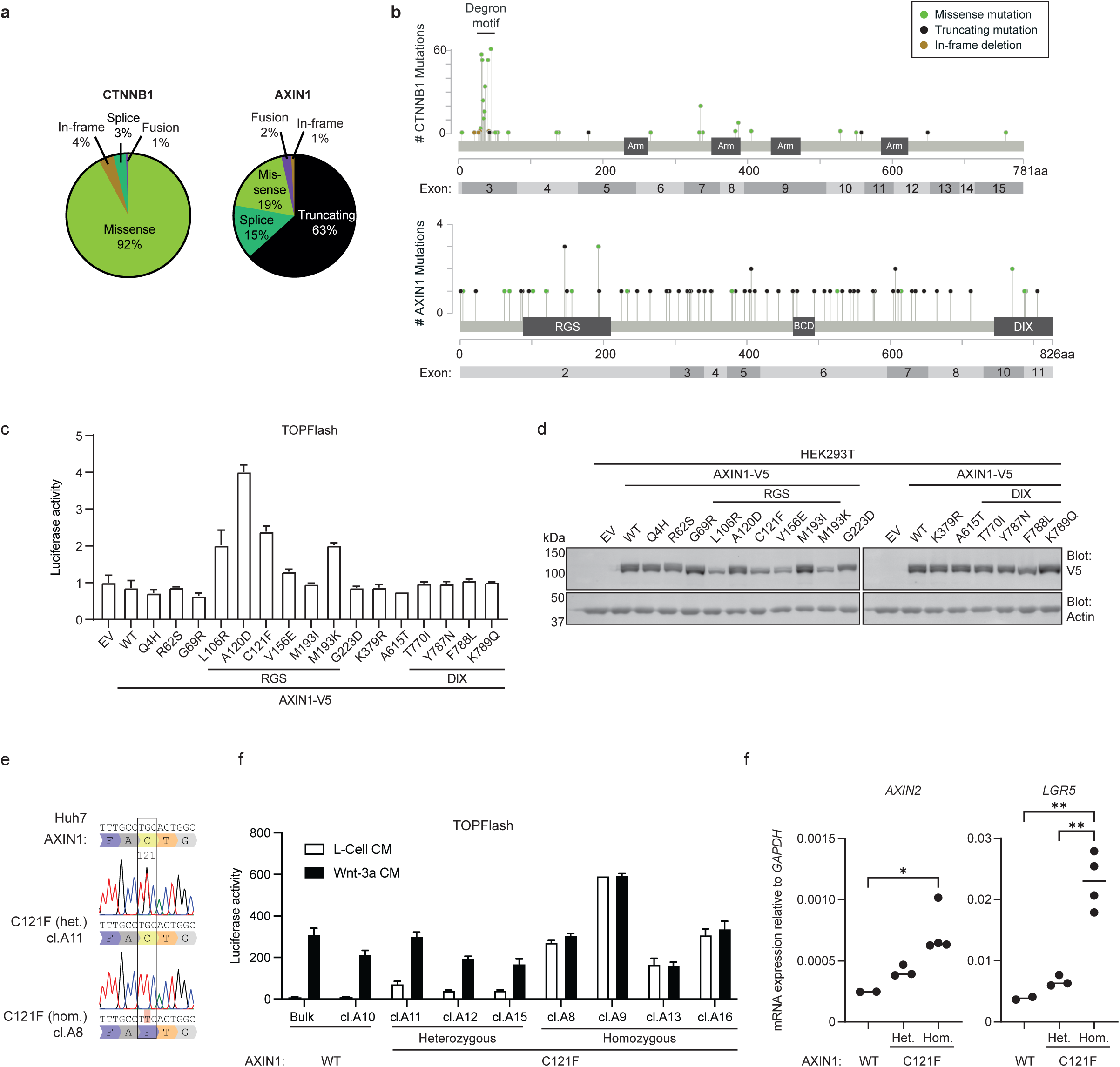
Missense mutations within the RGS domain of AXIN1 drive Wnt/β-catenin signaling. **(a)** Query of liver cancer data deposited by MERiC, INSERM, MSK, AMC, RIKEN and TCGA, containing sequencing data from a total of 1233 patient samples. **(b)** Schematic depicting the number of mutations per amino acid in *CTNNB1* and *AXIN1*, based on the same liver cancer databases as used in (a). Colour-coding depicts the type of mutation. Arm, Armadillo repeats; BCD, β-catenin binding domain. **(c)** TOPFlash reporter assay of overexpressed AXIN1 variants in HEK293T cells. EV, indicates empty vector control. Graph shows a representative experiment (N=3) with mean +/-SD of n=2 independent wells. Mutations in RGS or DIX domain are indicated. **(d)** Western blot of AXIN1-V5 expression levels of assay shown in (c). Actin was used as loading control. **(e)** Sanger sequencing of Huh7 clones harboring a heterozygous or homozygous base substitution in *AXIN1* exon 2, resulting in the C121F mutation. For all sequencing data, see figure S4. **(f)** TOPFlash reporter assay comparing non-modified Huh7 cells to clones harboring heterozygous and homozygous AXIN1 C121F mutations. Cells were treated with Wnt-3a conditioned medium or L-cell conditioned medium as control. Graph shows a representative experiment (N=3) with mean +/-SD of n=2 independent wells. **(g)** RT-qPCR experiments depicting expression of *AXIN1* and Wnt target genes *AXIN2* and *LGR5* relative to the household gene *GAPDH*. Each dot represents the mean of three biological replicates per clone. Significance was determined using one-way ANOVA analysis. * indicates p ≤ 0.05, ** indicates p ≤ 0.01. non-significant comparisons were left out for clarity.

To evaluate the functional consequences of RGS cancer mutations at the endogenous level, we introduced homozygous AXIN C121F and L106R mutations in HEK293T cells using prime editing and CRISPR/Cas9-assisted knock-in, respectively (Fig. S2a and c). All clones harboring these HCC-associated mutations displayed hyperactivation of Wnt/β-catenin signaling in the absence of Wnt supplementation (Fig. S2b and d). Furthermore, we confirmed loss of interaction between AXIN1 L106R and APC in these clones (fig. S2e-f), substantiating the previously observed loss of structural integrity of the RGS domain^15^. Together, these data suggest that cancer-associated AXIN1 RGS mutations promote Wnt/β-catenin signaling by disrupting destruction complex assembly and function.

Next, we examined how AXIN1 RGS missense cancer mutations impact Wnt/β-catenin signaling in a liver cancer background, using Huh7 cells. This cell line carries no mutations in core components of the Wnt/β-catenin pathway^29^, and is responsive to treatment with Wnt-3a and RSPO1, indicating that the Wnt/β-catenin signaling route is intact (Fig. S3). Furthermore, Huh7 cells were not sensitive to treatment with the Wnt secretion inhibitor LGK974, demonstrating that these cells do not depend on Wnt ligands for their growth and survival (Fig. S3). We employed prime editing to introduce AXIN1 C121F mutations in Huh7 cells (Fig. 1e; S4a)^30^. We obtained heterozygous and homozygous mutant clones, for which activation of Wnt/β-catenin signaling in the absence of Wnt ligand directly correlated with their mutant allele frequency (Fig. 1f). In line with these findings, expression of the Wnt target genes *AXIN2* and *LGR5* was elevated in a stepwise manner in heterozygous and homozygous AXIN1 C121F clones (Fig. 1g). Together, these data provide compelling evidence that AXIN1 RGS missense mutations induce Wnt/β- catenin pathway activation in liver cancer cells.

### Frameshift mutations in 5’ coding regions yield an N-terminally truncated AXIN1 variant with partially retained functionality

The majority of *AXIN1* mutations in HCC are frameshift mutations that, in case nonsense mediated decay is inefficient, may drive formation of truncated AXIN1 variants (Fig. 1a). Frameshift mutations that locate to 5’ regions of the coding parts of the gene, are predicted to lead to an AXIN1 deletion phenotype, either due to nonsense-mediated decay or loss of most functional domains in case of truncation^17,31^. In line with this assumption, Cre-mediated excision of *AXIN1* exon 2, encoding for amino acids 1 to 292, was used previously to generate conditional *AXIN1* knock-out mice^14^. We next aimed to assess the functional consequences of frameshift mutations in *AXIN1* close to the translation start site by using CRISPR/Cas9-mediated gene editing in Huh7 cells (Fig. 2a; S4b)^32^. In line with loss of AXIN1 suppressor function, these mutations induced basal Wnt/β-catenin signaling and the effects correlated with *AXIN1* mutant allele frequency (Fig. 2b). Unexpectedly, however, Western blotting revealed that these Huh7 clones retained expression of an AXIN1 fragment with a lower molecular weight, as shown by an antibody recognizing the central region of AXIN1 (AF3287) (Fig. 2c, lower arrow). This AXIN1 variant could also be detected using an antibody directed against the AXIN1 C-terminus (C76H11), indicating this concerned an N-terminally truncated variant (Fig. 2d, lower arrow). In corroboration, an overexpression construct encoding for mutation R22* also resulted in an AXIN1 variant with reduced molecular weight, which was detectable using a C-terminal V5 epitope tag, both on Western blot and by immunofluorescence (Fig. 2e, S5a). Furthermore, these N-terminally truncated AXIN1 variants maintained their capacity to form cytosolic puncta, confirming that their C-terminal DIX domain remained intact (Fig. S5a). Truncations resulting from frameshift mutations downstream of AXIN1 position R146 could not be visualized via the C-terminal V5 tag, while these variants were still detected by the AF3287 antibody, recognizing the central part of AXIN1 (Fig. 2e, S5a). Thus, while stop codons downstream of R146 generate C-terminally truncated AXIN1 variants, introduction of stop codons in the 5’ region of *AXIN1* exon 2 result in AXIN1 truncations that lack the N-terminus (ΔN) but still harbor the C-terminal DIX domain. Based on these findings, we hypothesized that ΔN AXIN1 variants may be generated by alternative translation initiation^33^. To examine this assumption, we eliminated potential alternative start sites by mutating methionines M173, M181, M193 and M253 in exon 2 to leucine within AXIN1 R22* variants (Fig. 2f). Indeed, combined mutation of M173 and M253 disrupted expression of the AXIN1 R22* variant, indicating that these start sites are utilized for alternative translation initiation (Fig. 2g).

**Figure 2.**
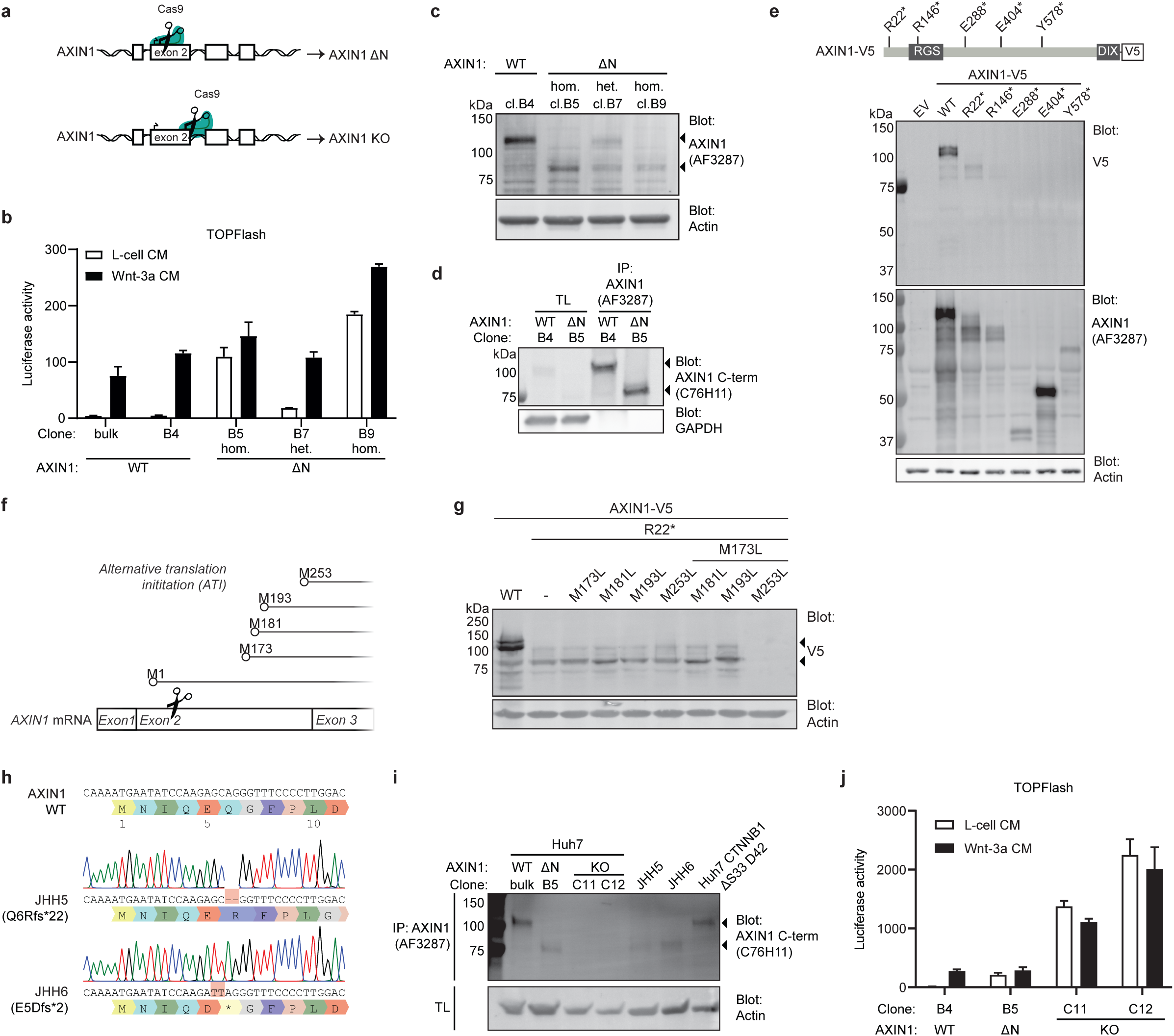
Frameshift mutations in 5’ coding regions yield an N-terminally truncated AXIN1 variant with partially retained functionality. **(a)** Schematic depicting the different single guide RNA recognition sites used to generate CRISPR/Cas9-mediated frameshifts in the *AXIN1* locus. Targeting close to the translation initiation site results in an N-terminally truncated (ΔN) AXIN1 variant, whereas targeting in the 3’ region of exon 2 results in a full AXIN1 knock-out (KO). **(b)** TOPFlash reporter assay comparing non-modified Huh7 cells to clones with heterozygous and homozygous frameshift mutations early in the *AXIN1* gene. Huh7 cells were treated with Wnt-3a conditioned medium or L-cell conditioned medium as control. Graph shows a representative experiment (N=3) with mean +/-SD for n=2 independent wells. **(c)** Western blot of endogenous AXIN1 levels in Huh7 clones harboring *AXIN1* frameshift mutations close to the translation start site. Arrows highlight the AXIN1 WT and ΔN variants. Actin was used as loading control. **(d)** Western blot of immunoprecipitated AXIN1 from Huh7 WT clone B4 and AXIN1 ΔN clone B5. Immunoprecipitation was performed to enrich for AXIN1 for visualization purpose. **(e)** Top: Schematic depicting AXIN1 with a C-terminal V5 tag. Lines indicate the sites at which stop codons are introduced. Termination mutations are all derived from the HCC databases described in figure 1b. Bottom: Western blot of HEK293T lysates overexpressing AXIN1-V5 truncated variants stained for V5 and AXIN1. Actin was used as loading control. EV, empty control vector. **(f)** Schematic depicting candidate start sites for recognition by the translation initiation machinery upstream in the *AXIN1* locus. The 4 methionines found in the 3’ part of exon 2 are the main candidates for alternative translation initiation. **(g)** Western blot of HEK293T cells overexpressing AXIN1 variants harboring truncating mutation R22* in combination with different methionine-to-leucine substitutions (similar side chain size), stained for V5 and AXIN1. Actin was used as loading control. **(h)** Sanger sequencing of HCC cell lines JHH5 and JHH6, both harboring 5’ frameshift mutations in *AXIN1*. **(i)** Western blot of immunoprecipitated AXIN1 from Huh7, JHH5, JHH6 and different *AXIN1*-mutated Huh7 cell lines. Actin was used as loading control. **(j)** TOPFlash reporter assay comparing non-modified Huh7 cells to clones with different AXIN1 truncations. Cells were treated with Wnt-3a conditioned medium or L-cell conditioned medium as control. Graph shows a representative experiment (N=3) with mean +/- SD of n=2 independent wells.

To test whether alternative translation initiation is physiologically relevant in the context of liver cancer, we evaluated AXIN1 expression in the HCC cell lines JHH5 and JHH6 that both harbor 5’ truncating mutations in exon 2 (Q6Rfs*22 and E5Dfs*2, respectively) (Fig. 2h). Indeed, these cell lines expressed an AXIN1 variant at a similar molecular weight as our CRISPR/Cas9-modified Huh7 cells with 5’ truncations in *AXIN1* exon 2 (Fig. 2i). Importantly, due to these alternative translation initiation sites, a robust AXIN1 knock-out (KO) line would require truncating mutations downstream of the initiation sites M173 and M253. Indeed, introduction of frameshift mutations immediately downstream of M253 led to loss of detectable AXIN1 expression (Fig 2i; S4c). Of note, Huh7 AXIN1 KO cells displayed strongly increased levels of basal Wnt/β-catenin pathway activity in comparison to Huh7 AXIN1 ΔN cells (Fig. 2j). In agreement, AXIN1 KO HEK293T cells also displayed increased Wnt/β-catenin signaling compared to AXIN1 ΔN HEK293T cells (Fig. S5b-d). Together, these results reveal a previously overlooked class of *AXIN1* 5’ frameshift mutations that drive expression of an N-terminally truncated protein with partially retained functionality.

### *AXIN1*-mutant HCC cells display moderate Wnt/β-catenin signaling, while *CTNNB1*-mutant cells are Wnt-high

While activating *CTNNB1* mutations have been associated with Wnt/β-catenin hyperactivation in HCC, *AXIN1* mutations were linked to low or even completely absent Wnt/β-catenin signaling^10,13^. To directly compare both mutation types in Huh7 cells, we introduced an in-frame deletion of β-catenin degron residue S33 using prime editing (Fig. S4d)^34^. Clones carrying homozygous β-catenin ΔS33 mutations displayed higher Wnt/β-catenin pathway activation compared to different classes of AXIN1-mutant clones (Fig. 3a). At the transcriptional level, *CTNNB1*-mutant Huh7 cells displayed increased expression of Wnt target genes *AXIN2* and *LGR5* (Fig. 3b). In line with the luciferase reporter data, these genes were upregulated to a lower extent in *AXIN1*-mutant lines (Fig. 3b). In further agreement, β-catenin levels were more strongly stabilized in *CTNNB1*-mutant versus WT and *AXIN1*-mutant Huh7 cells (Fig. 3c-d). No significant difference in β-catenin accumulation between WT and *AXIN1*-mutated Huh7 clones was observed, corroborating the notion that β-catenin is an insensitive marker for Wnt pathway activation^17^. Next, we queried the TCGA liver carcinoma database to examine the expression of *AXIN2* and *LGR5* in HCC patient samples (Fig. 3e). *AXIN2* expression was slightly (but not significantly) upregulated in *AXIN1*-mutant HCC, while this Wnt target gene was significantly elevated in *CTNNB1*-mutant HCC. In agreement with our observations in Huh7 cells, *LGR5* expression displayed a significant upregulation in *AXIN1*-mutant HCC and these effects were more pronounced in *CTNNB1*-mutant HCC samples. Together, our results show that both *AXIN1*- mutant and *CTNNB1*-mutant HCC display elevated Wnt/β-catenin signaling, although levels of pathway activation are more moderate for *AXIN1*-mutant cells.

**Figure 3.**
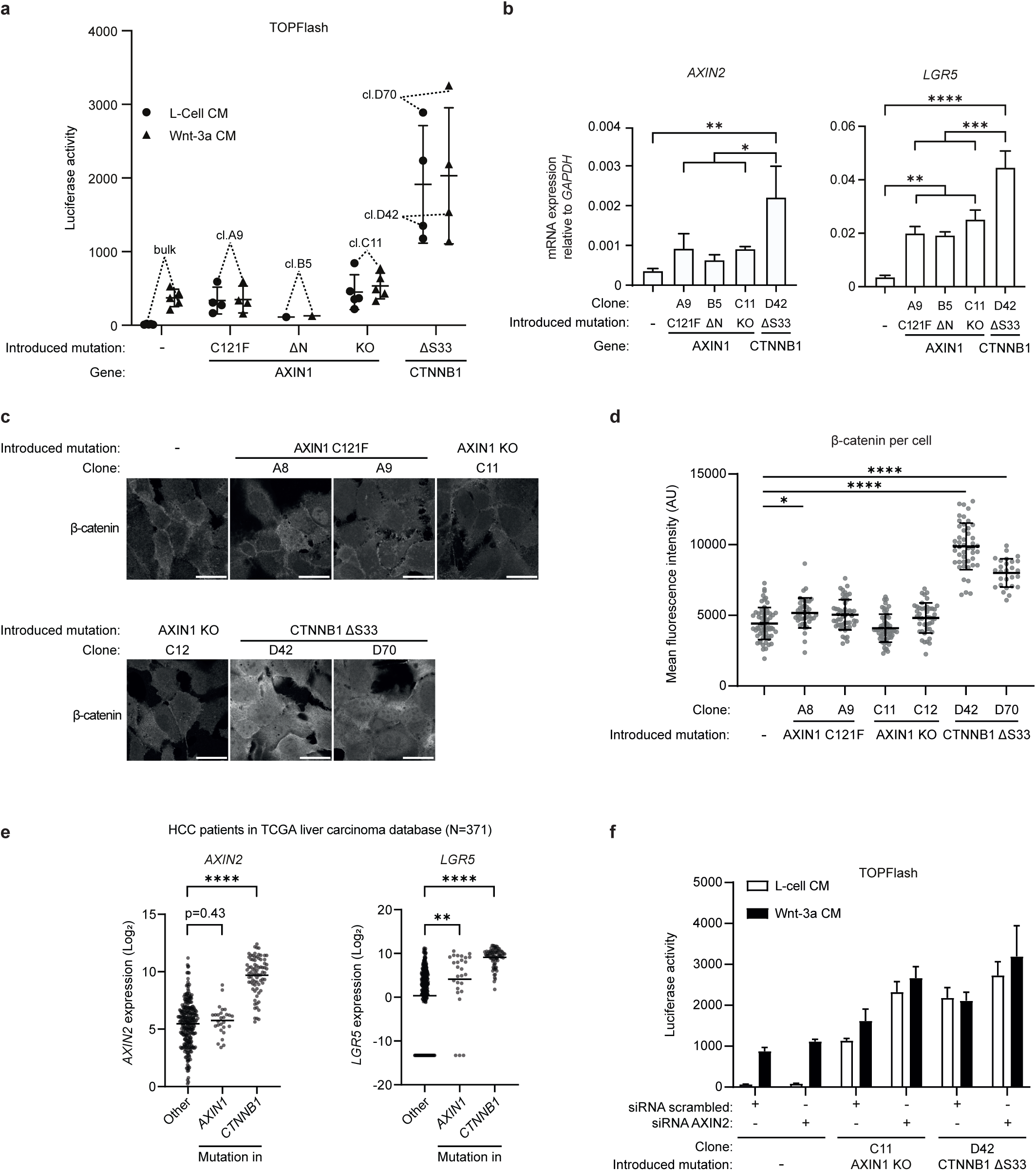
*AXIN1*-mutant HCC cells display moderate Wnt signaling levels, while *CTNNB1*- mutant cells are Wnt-high. **(a)** TOPFlash reporter assay comparing non-modified Huh7 cells to clones carrying different *AXIN1* and *CTNNB1* mutations. Graph shows a representative experiment (N=3), where one dot represents the mean of technical duplicates of one clone. Cells were treated with Wnt-3a conditioned medium or L-cell conditioned medium as control. **(b)** RT- qPCR depicting expression of Wnt target genes *AXIN2* and *LGR5* relative to the household gene *GAPDH*. Graph shows the mean of three biological replicates per clone +/- SD. Significance was determined using one-way ANOVA. **(c)** Immunofluorescence images of Huh7 lines harboring different Wnt pathway mutations, labelled for β-catenin. Scale bar represents 30 µm. **(d)** Quantification of immunofluorescence images belonging to the experiment in (c). Quantified using an automated ImageJ script for which details are described in the methods section. Significance was determined using one-way ANOVA. **(e)** Analysis of expression data from the TCGA liver carcinoma database, accessed using the R2 analysis platform. Since log_2_ values cannot equal 0, all missing values in this dataset have been replaced with 0.0001, of which the log_2_ equals - 13.2877. Significance was determined using one-way ANOVA. **(f)** TOPFlash reporter assay comparing non-modified Huh7 cells to clones harboring *AXIN1* and *CTNNB1* mutations after siRNA-mediated knock-down of *AXIN2*. * indicates p ≤ 0.05, ** indicates p ≤ 0.01, *** indicates p ≤ 0.001, **** indicates p ≤ 0.0001. Non-significant comparisons were left out for clarity.

AXIN2 normally acts as safeguard that curbs Wnt signaling in the absence of AXIN1^17,35,36^. By contrast, β-catenin mutants with a disrupted degron motif are insensitive to AXIN2 levels^35^. This may explain the difference in Wnt pathway activity between *CTNNB1*- and *AXIN1*-mutant HCC. Indeed, *AXIN2* knock-down in *AXIN1*-mutant Huh7 cells resulted in an increase in Wnt/β- catenin pathway activity to similar levels as *CTNNB1*-mutant cells (Fig. 3f). Similarly, CRISPR/Cas9-mediated inactivation of *AXIN2* in *AXIN1*-mutant HEK293T cells resulted in synergistic Wnt/β-catenin pathway activation (Fig. S6). These data confirm that elevated levels of AXIN2 partially compensate for impaired activity of its paralogue in *AXIN1*-mutant tumors.

### Wnt/β-catenin pathway activation leads to dose-dependent inhibition of YAP/TAZ signaling

Previous studies have linked the occurrence of *AXIN1* mutations in HCC to inactivation of the Hippo signaling pathway and concomitant hyperactivity of YAP/TAZ-mediated transcription, although the underlying mechanisms have remained incompletely understood^10,11^. To address this issue, we set out to compare the levels of YAP/TAZ signaling in Huh7 cells in the absence or presence of mutations in *AXIN1* and *CTNNB1*. Since the Hippo signaling cascade is primarily regulated by mechanotransduction, we first assessed whether this pathway is sensitive to alterations in Huh7 cell confluency^37^. Compared to Huh7 cells at lower densities, cells that were growing near confluency displayed nuclear exclusion of YAP (Fig. 4a), decreased YAP/TAZ-mediated luciferase reporter activity (Fig. 4b) and downregulated expression of YAP/TAZ targets *CYR61*, *CTGF*, *AREG* and *ANKRD1* (Fig. 4c). These data indicate that the Hippo pathway is functional in Huh7 cells.

**Figure 4.**
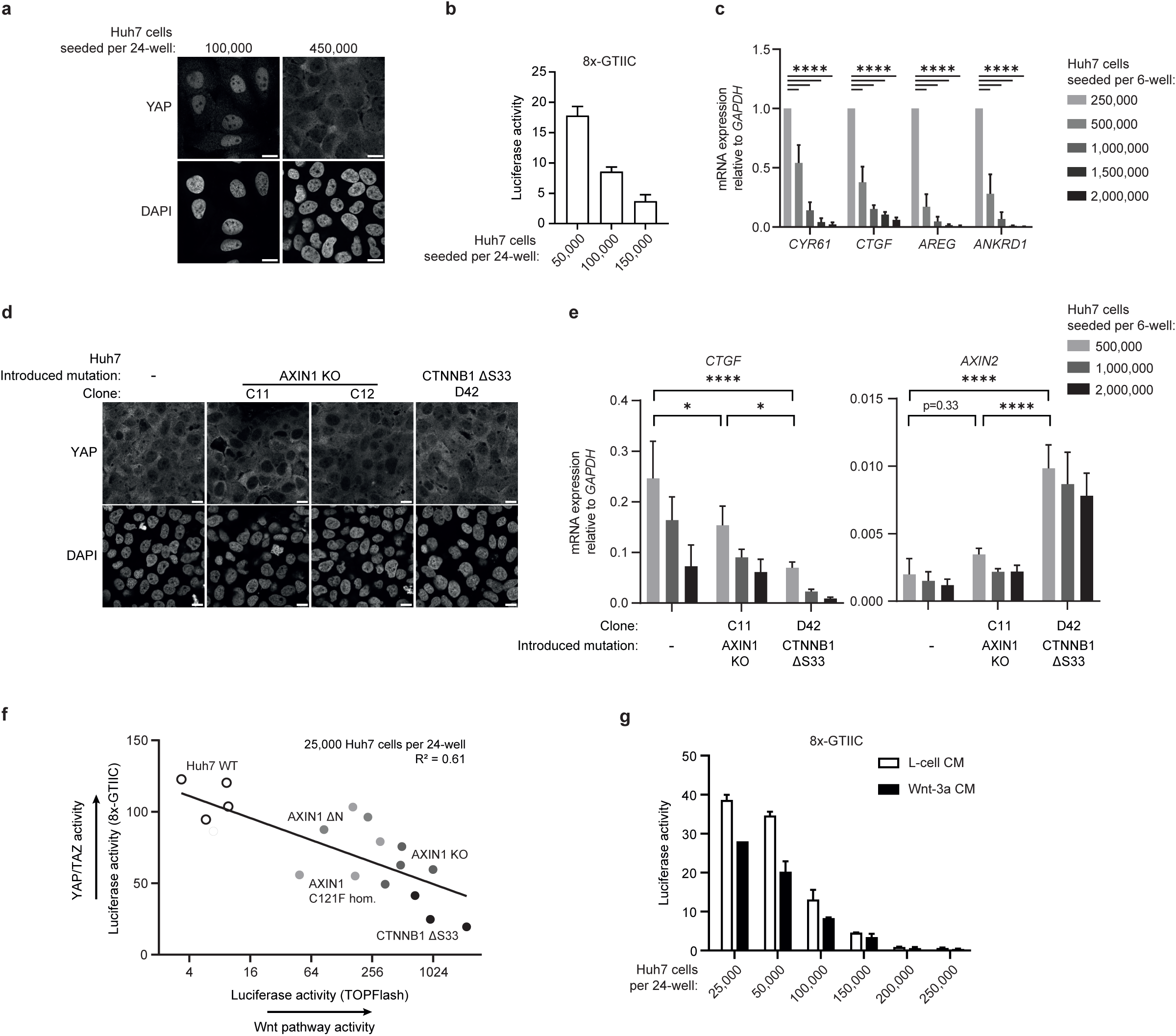
Wnt pathway activation leads to dose-dependent inhibition of YAP/TAZ signaling. **(a)** Representative immunofluorescence images of Huh7 cells seeded at low and high cell density. Fixed cells were labeled for YAP and DAPI. Scale bar represents 30 µm. **(b)** 8xGTIIC reporter assay comparing Huh7 cells seeded at different cell densities. Graph shows a representative experiment with mean +/- SD of n=2 independent wells. **(c)** RT-qPCR depicting expression of YAP/TAZ target genes *Cyr61*, *CTGF*, *AREG* and *ANKRD1* relative to the household gene *GAPDH* in Huh7 cells seeded at different cell densities. Each condition was normalized to the condition with the lowest seeding density. Bars and error bars represent the mean of three biological replicates +/- SD. Significance was determined using two-way ANOVA. **(d)** Representative immunofluorescence images of Huh7 clones with different genotypes seeded at 200,000 cells per 24-well. Fixed cells were labeled for YAP and DAPI. Scale bar represents 30 µm. **(e)** RT-qPCR depicting expression of YAP/TAZ target gene *CTGF* and Wnt target gene *AXIN2* relative to the household gene *GAPDH* for different Huh7 clones at increasing cell density. To directly compare all clones, relative quantification (RQ) values representing the expression compared to *GAPDH* were plotted. Bars and error bars represent the mean and standard deviation of three biological replicates. Significance was determined using two-way ANOVA. **(f)** Scatter plot comparing YAP-responsive reporter activity to TOPFlash reporter activity for a set of 17 Huh7 clones with different genotypes. Representative of N=3 experiments. The coefficient of determination (R2) was determined using GraphPad analysis software. **(g)** 8xGTIIC luciferase assay comparing Huh7 cells seeded at different cell densities, either treated with Wnt-3a conditioned medium or L-cell medium as control. Graph shows a representative experiment (N=3) with mean and standard deviation of duplicate cell cultures transfected in parallel. * indicates p ≤ 0.05, **** indicates p ≤ 0.0001. Non-significant comparisons were left out for clarity.

We next assessed whether Wnt/β-catenin pathway alterations affect YAP localization in Huh7 cells seeded at relatively high confluency. Nuclear YAP levels were not visibly altered in *AXIN1*-deficient Huh7 cells compared to non-modified control cells (Fig. 4d, S7a). Moreover, both *AXIN1* and *CTNNB1*-mutant cells displayed reduced levels of YAP/TAZ-mediated transcription when compared with control cells at varying seeding densities (Fig. 4e; S7b). To perform a more comprehensive investigation of the relationship between Wnt/β-catenin pathway activity and Hippo signaling, we examined how varying levels of β-catenin-mediated transcription affect YAP/TAZ-dependent reporter activities in a large set of *AXIN1*-or *CTNNB1*-mutant Huh7 clones (Fig. 4f). The results clearly revealed an inverse correlation between levels of Wnt/β-catenin pathway activation and YAP/TAZ transcriptional activity (Fig. 4f). In an earlier study, the liver-specific Wnt target gene TBX3 was reported to repress YAP/TAZ signaling, providing a potential mechanism for the negative correlation between both pathways^38^. We however did not detect upregulated levels of TBX3 in either AXIN1- or CTNNB1-mutant cells, excluding this possibility as an explanation of our results (Fig. S7c). To further investigate how both pathways are interlinked, we treated control Huh7 cells with Wnt-3a conditioned medium (Fig. 4g). Wnt-3a supplementation negatively regulated YAP/TAZ activation in a reporter assay, suggesting that there is crosstalk between both pathways that is not exclusive to *AXIN1*- or *CTNNB1*-mutant Huh7 cells. Next, we used different small molecules to modulate Wnt/β-catenin and YAP pathway activity. Notably, treatment with the small molecule K-975, which inhibits TEAD transcription by blocking its interaction with YAP/TAZ, did not affect Wnt/β-catenin signaling in any of the Huh7-derived cell lines (Fig. S7d-f), suggesting that alterations in YAP/TAZ-mediated transcription do not affect Wnt/β-catenin pathway activity. On the other hand, treatment with CHIR99021, a GSK3 inhibitor that activates Wnt/β-catenin signaling, downregulated expression of YAP target genes *CTGF, CYR61 and ANKRD1* in both non-modified and *AXIN1*-mutant Huh7 cells (Fig. S7d-e). Importantly, similar effects of CHIR99021 treatment were observed in *CTNNB1*-mutant cells, indicating that GSK3 inhibition might act on YAP/TAZ signaling by a mechanism partially independent of β-catenin-dependent transcription (Fig. S7f). Together, our results indicate that Wnt pathway activation negatively impacts transcription of YAP/TAZ-associated target genes in a dose-dependent manner in hepatocellular carcinoma cells.

### *Axin1* mutations promote Wnt ligand-independent growth in mouse liver progenitor organoids

Our results argue that *AXIN1* mutations drive Wnt/β-catenin signaling in HCC (Fig. 3a). To determine whether these mutations suffice to acquire niche factor independence in a physiological setting, we introduced Axin1-truncating mutations in a mouse liver progenitor organoid line (Fig. S8a-b)^39^. In line with our findings in human cells, the introduction of frameshift mutations in upstream 5’ regions (residue 15) led to the expression of N-terminally truncated Axin1 variants (Axin1 ΔN), whereas frameshift mutations introduced more downstream (residue 287) induced loss of detectable Axin1 protein (Axin1 KO) (Fig. 5a; S8c). Proliferation and propagation of mouse liver progenitor organoids largely depends on Wnt/β-catenin pathway activation by RSPO1 supplementation and autocrine Wnt secretion^39^. Strikingly, both Axin1 ΔN and Axin1 KO liver organoids displayed RSPO1-independent growth and were insensitive to treatment with the Wnt secretion inhibitor LGK974 (Fig. 5b-d; S8d-e)^40^. Notably, we observed similar expansion rates for both Axin1 ΔN and Axin1 KO lines upon RSPO1 withdrawal and porcupine inhibitor treatment, suggesting that both classes of *Axin1* mutations can drive niche factor-independence in the liver (Fig. 5d).

**Figure 5.**
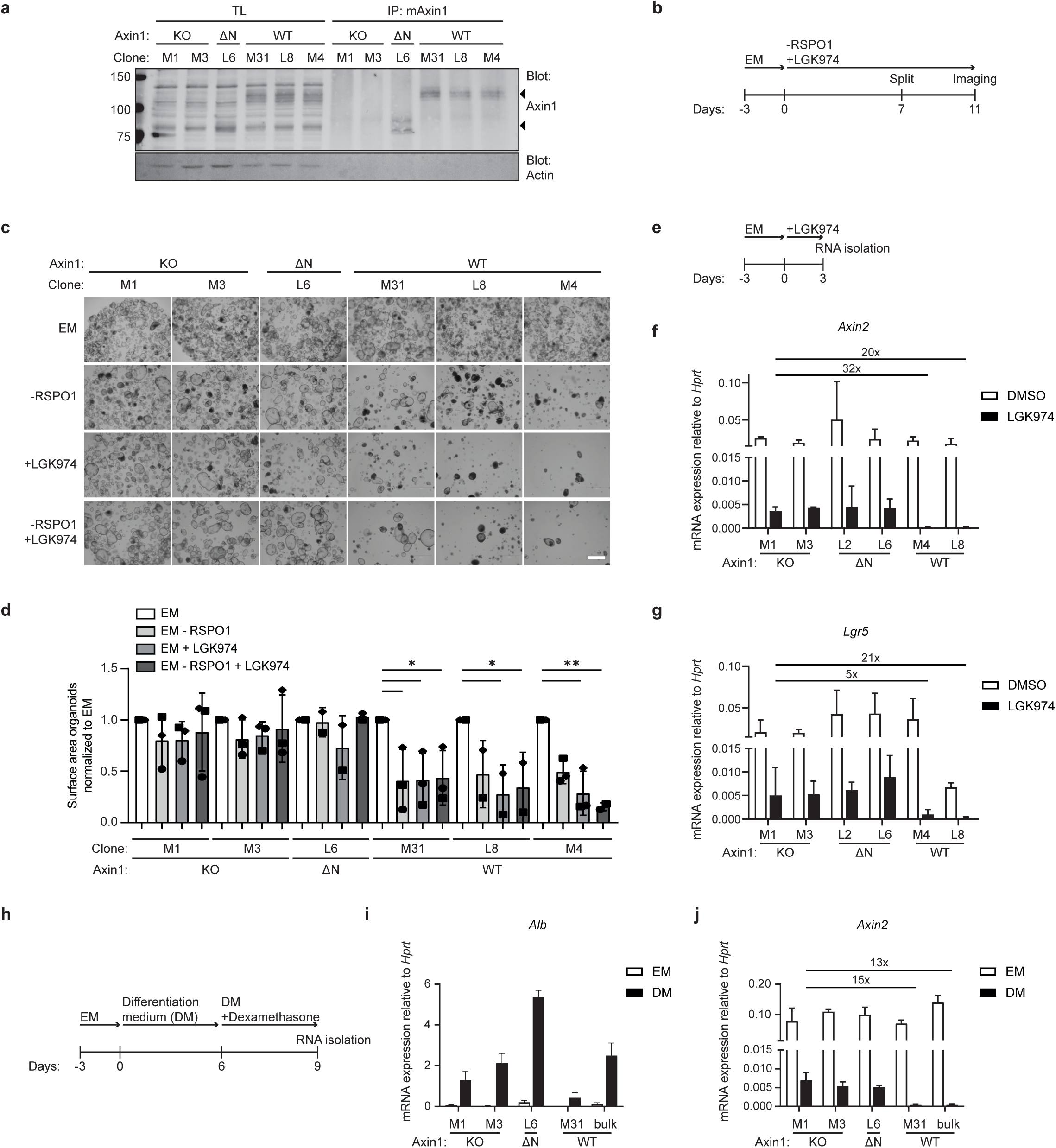
*Axin1* mutations promote Wnt ligand-independent growth in mouse liver progenitor organoids. **(a)** Western blot of immunoprecipitated Axin1 from WT and *Axin1*-mutant mouse liver progenitor organoids. Actin was used as loading control. **(b)** Diagram displaying the setup of the cell viability experiment performed (c) and (d). **(c)** Representative brightfield images of liver progenitor organoids cultured in expansion medium (EM) with and without 10% RSPO1 conditioned medium and 500 nM LGK974. Scale bar, represents 500 µm. EM, expansion medium. **(d)** Quantification of biological replicates as performed in (c). Quantification was performed by determining cell surface area of the organoids using OrganoSeg analysis software^61^. Each condition was normalized to organoids cultured in EM. EM, expansion medium. * indicates p ≤ 0.05, ** indicates p ≤ 0.01. **(e)** Diagram of the experiment setup for the RT-qPCR depicting expression of **(f)** *Axin2* and **(g)** *Lgr5* relative to household gene *Hprt*. Organoids were treated with 500 nM LGK974 for 3 days. Bars and error bars represent the mean of three biological replicates +/- SD. Fold-change differences between the indicated conditions were shown. **(h)** Diagram displaying the differentiation protocol. **(i-j)** RT-qPCR experiments for *Alb* (coding for Albumin) and *Axin2* mRNA levels, respectively. Fold-change differences between the indicated conditions were depicted. Bars and error bars represent the mean of three biological replicates +/- SD. EM, expansion medium; DM, differentiation medium.

To determine how *Axin1* mutations influence Wnt/β-catenin pathway activity in these organoids, we assessed the effect of short-term LGK974 treatment on Wnt target gene transcription (Fig. 5e-g). Although treatment drastically reduced expression of *Lgr5* and *Axin2* in both WT and mutant organoid lines, residual expression of these Wnt target genes was approximately 5- to 40-fold times higher in both *Axin1*-mutant lines (Fig. 5f-g). Of note, LGK974 treatment did not affect *Axin1* expression (Fig. S8f). Together, these results suggest that low levels of Wnt/β-catenin signaling are sufficient to sustain proliferation of liver progenitor cells. To determine the effects of *Axin1* mutations in organoids with hepatocyte features, we applied a previously defined differentiation protocol (Fig. 5h)^39^. All organoid lines were capable of differentiating towards the hepatocyte lineage in these conditions, as evidenced by increased expression of Albumin (Fig. 5i). Despite their induced differentiation, *Axin1*-mutant organoids retained a 15-fold increase in expression of the Wnt target gene *Axin2* when compared to WT organoid lines (Fig. 5j). Together, these results indicate that *Axin1* mutations promote Wnt/β- catenin pathway activity in organoids with liver progenitor as well as hepatocyte characteristics.

## Discussion

Mutations in the Wnt/β-catenin pathway-associated genes *AXIN1* and *CTNNB1* are inherently linked to the development of HCC^2,41^. Recent genome sequencing efforts of HCC patient cohorts revealed that mutations in *AXIN1* and *CTNNB1* are mutually exclusive and associate with distinct tumor characteristics, aggressiveness and clinical prognosis^9^. In this study, we aimed to examine the differences in downstream signaling events of each mutation type in HCC. Consistent with earlier studies, Wnt/β-catenin pathway activation was strongly induced upon introduction of β-catenin mutations in the liver cancer cell line Huh7^13,42,43^. By contrast, introduction of *AXIN1* mutations induced only low levels of Wnt/β-catenin signaling, but with clear physiological relevance, as illustrated by the acquired Wnt/RSPO1-independent growth properties of *AXIN1*- mutant mouse liver progenitor organoids. Importantly, our findings therefore challenge the claim that *AXIN1* mutations drive HCC development independent of the Wnt/β-catenin pathway^10^. Notably, *Axin1*-mutant organoids that were treated with the Wnt secretion inhibitor LGK974 still displayed a reduction in Wnt target gene expression, suggesting that *AXIN1* mutations do not lead to complete uncoupling of the signaling cascade from the Wnt receptor complex. Thus, *AXIN1* mutations likely provide a low level of Wnt/β-catenin signaling that is sufficient for survival and proliferation, and still allows for regulation by upstream signaling events. These findings are reminiscent of an earlier report describing that residual levels of MAPK signaling in colon organoids carrying a KRAS mutation were sufficient to evade apoptosis in the absence of EGF ligand^44^. In line with our findings, optimal Wnt signaling levels for HCC formation were estimated to be lower than those required for intestinal tumors, further supporting the view that *AXIN1* mutations facilitate ‘just-right’ levels of Wnt signaling to enable HCC development^45,46^.

The majority of AXIN1 mutations found in HCC are truncating and missense mutations. Although we and others have previously established a Wnt pathway-promoting role for missense mutations in the AXIN1 RGS domain, the physiological relevance of this mutation class in liver cancer has not been determined^15–17^. Here, we confirm that RGS-associated AXIN1 missense mutations are capable of driving Wnt/β-catenin signaling in liver cancer cells. In addition, we found that truncating mutations near the canonical start site of *AXIN1* result in alternative translation initiation and, consequently, expression of an N-terminally truncated variant with residual tumor suppressor function. Notably, alternative translation initiation may represent a wider mechanism for tumor suppressors in cancer, considering a report claiming that generally more than 50% of CRISPR-cas9-mediated “knock-out” cell lines retain expression of the targeted gene^33^. As AXIN1 ΔN variants lack the tankyrase (TNKS)-binding and RGS domains, tumors harboring these mutations will be insensitive to TNKS1/2 inhibitors that stabilize AXIN1 or to the small molecule KYA1797, that binds AXIN1 RGS and increases β-catenin phosphorylation by promoting GSK3 activity^47,48^. Instead, inhibition of SIAH1/2, ubiquitin ligases that regulate AXIN1 stability by associating with its more downstream GSK3 binding domain, might provide an effective strategy to stabilize AXIN1 ΔN cancer variants^49^. Together, our findings underscore the need for careful evaluation of different cancer mutation types to improve patient stratification^7^.

Several studies have linked AXIN1 deficiency to increased signaling of the Hippo pathway effectors YAP and TAZ^10,11,50^. Strikingly, however, none of the AXIN1 variants that we studied induced YAP/TAZ signaling in Huh7 cells, in agreement with a recent study^16^. By contrast, elevated Wnt/β-catenin pathway activation attenuated YAP/TAZ signaling in a dose-dependent manner. These observations are in line with several studies that reported a negative correlation between both pathways in the liver^51,52^. Our results suggest that the low levels of Wnt pathway activation induced by *AXIN1* mutations are sufficient to drive proliferation while, at the same time, providing a permissive environment for moderate levels of YAP/TAZ signaling. These effects of *AXIN1* mutations stand in contrast with the strong levels of Wnt/β-catenin pathway activation induced by *CTNNB1* mutations, which drive suppression of YAP/TAZ signaling. As mutations in core Wnt/β-catenin pathway components are late events in HCC development and *Axin1*-deficient mice show delayed tumorigenesis (>1 year)^10,14,53^, we anticipate that additional mutations are necessary for HCC progression and that differences in mutational background may impact on the degree of YAP/TAZ activation in these cancer cells. In addition, HCC is often associated with alterations in the tumor microenvironment, including stiffening of the extracellular matrix, which directly links to increased YAP/TAZ signaling^54^. These factors might explain why transcriptional hyperactivation of YAP/TAZ in *Axin1*-deficient mouse tumors is not observed in our *AXIN1*-mutant cell cultures^10,11^.

In conclusion, our study reveals that different classes of *AXIN1* mutations endow liver cells with physiologically relevant Wnt/β-catenin signaling levels, although pathway activation is less pronounced than observed for *CTNNB1* mutations. Our results however indicate that the levels of Wnt signaling induced in *AXIN1*-mutant liver organoids is sufficient to sustain growth in the absence of Wnt ligands, explaining why Wnt pathway inhibition in *AXIN1*-mutant HCC has so far proven unsuccessful^35^. We propose that combination treatment with inhibitors for Wnt/β-catenin and YAP/TAZ signaling may offer a promising therapeutic avenue for HCC tumors harboring *AXIN1* mutations^55,56^.

## Methods

### Cell culture

Human embryonic kidney (HEK293T) cells were cultured in Dulbecco’s modified Eagle’s medium with 4500 mg/L glucose (Sigma-Aldrich). Huh7 cells were cultured in Dulbecco’s modified Eagle’s medium with 1000 mg/L glucose (Sigma-Aldrich). JHH5 and JHH6 cells were cultured in William’s E medium (Gibco). All cell culture media were supplemented with 10% fetal bovine serum (GE Healthcare), 2 mM L-alanyl-L-glutamine (Sigma-Aldrich), 100 units/mL penicillin and 100 μg/mL streptomycin (Invitrogen). Cells were cultured in 5% CO2 at 37 °C. Wnt-3a conditioned medium was obtained from mouse L-cells stably expressing and secreting mouse Wnt-3a. L-cell conditioned medium was obtained from control L-cells. RSPO1 conditioned medium was produced using HEK293T cells stably transfected with human RSPO1-V5. Wnt surrogate conditioned medium was produced by conditioned medium of HEK293T cells transfected with Wnt surrogate (kind gift from Claudia Janda, Prinses Maxima Center, Netherlands)^57^ and added to cultures at 0.5-1%.

### Organoid culture

C57BL/6 mouse liver progenitor organoids were isolated as previously described^39^. For culturing, organoids were mixed with 40 µL matrigel (BD bioscience) per droplet and seeded in a 24-well plate and grown in expansion medium (EM). Expansion medium was based on Advanced DMEM F-12 (Gibco) supplemented with 100 units/mL penicillin and 100 μg/mL streptomycin (Invitrogen), 2 mM L-alanyl-L-glutamine (Sigma-Aldrich), 10 mM HEPES (Gibco), 1% B27 (Life Technologies), 1.25 μM N-acetylcysteine (Sigma-Aldrich), 10 nM gastrin (Sigma-Aldrich), 50 ng/mL mouse Egf (Peprotech), 10% RSPO1-V5 conditioned medium, 100 ng/mL mouse Fgf10 (Peprotech), 10 mM nicotinamide (Sigma-Aldrich) and 25 ng/mL mouse Hgf (Peprotech). Organoids were split in a 1:3- 1:6 ratio once every week by mechanical disruption. For differentiation, RSPO1-V5 condition medium, Hgf and nicotinamide were removed from the expansion medium, and instead A8301 (50 nM, Tocris) and DAPT (10 nM, Sigma) were added. The last 3 days of differentiation, dexamethason (30 µM, Sigma-Aldrich) was supplemented to the medium. For growth dependency assays, organoid lines were split 1:4 and directly cultured in EM, EM without RSPO1, EM supplemented with 500 nM LGK-974 (Cayman chemicals), or EM without RSPO1 and supplemented with 500 nM LGK-974. Organoids were split 1:4 after 7 days of culturing and were grown for 5 more days. Culture medium was replaced every other day. For short LGK-974 treatment, organoid lines were split 1:4 and cultured for 3 days with or without supplementation of 500 nM LGK-974.

### Constructs

TOPFlash and FOPflash luciferase reporter plasmids and thymidine kinase (TK)-Renilla were described previously ^58^. 8xGTIIC luciferase reporter was a kind gift from the lab of Martijn Gloerich (University Medical Center Utrecht, Netherlands), which is modified from a previously described vector^59^. For AXIN1 overexpression, human *AXIN1* isoform b was cloned into pcDNA3.1+ containing a C-terminal V5 tag. *AXIN1* mutations were introduced by site-directed mutagenesis using Q5 high-fidelity polymerase (New England Biolabs) using primers produced by Integrated DNA Technologies (see table S1). The sgRNA CRISPR/Cas9 vector PX459 V2.0 was kindly provided by the lab of Feng Zhang (Addgene, # 62988). Single guide RNAs (sgRNA) were cloned into the PX459 plasmid according to protocol^32^ (See Table S1 for guide sequences including overhangs). Prime editing guide RNAs were cloned into the pU6-pegRNA-GG-Vector, kindly provided by the lab of David Liu (Addgene, #132777). Annealed pegRNA spacers, pegRNA extensions, and pegRNA scaffold sequences with appropriate overhangs were inserted into the pU6-pegRNA-GG-Vector using Golden Gate assembly using BsaI-HfV2 (New England Biolabs) and T4 DNA ligase (New England Biolabs)^30^. For PE3 cloning, phU6-gRNA vector was digested using BbsI (Thermo Fisher) and subsequently ligated by T7 DNA ligase (New England Biolabs). CTNNB1 ΔS33 pegRNA and PE3 constructs were a kind gift from the lab of Sabine Fuchs^34^. A modified version of the PE2 plasmid by the lab of David Liu (Addgene, #132775) containing PE2 coupled via a P2A peptide to a puromycin resistance cassette was kindly provided by the lab of Susanne Lens (University Medical Center Utrecht, Netherlands).

### CRISPR/Cas9 and prime editing of cell lines

Huh7 and HEK23T cells were seeded in 10 cm dishes at 25% cell confluency and transfected one day later. For conventional CRISPR-Cas9-mediated targeting, cells were transfected using Fugene6 (Promega), according to the manufacturer’s protocol, with a DNA mixture of 6 μg PX459 plasmid containing sgRNA and 1 μg CMV-GFP plasmid to validate transfection efficiency. For prime editing, Huh7 cells were transfected using Fugene6 (Promega), with a DNA mixture containing 7.8 μg PE2-P2A-Puromycin plasmid, 2.6 μg pegRNA plasmid, 0.84 μg nicking sgRNA plasmid and 1 μg CMV-GFP. For both prime editing and CRISPR/Cas9 targeting, puromycin selection was started after 2 days by the addition of 2 μg/mL puromycin and maintained for an additional two days. After outgrowth, genomic DNA was isolated from the bulk population using the QiaAmp DNA micro kit (Qiagen) according to protocol. Genotyping of the bulk population was performed by genomic PCR using primers in Table S1 with either GoTaq Flexi DNA polymerase (Promega), or Q5 polymerase (NEB). PCR products were isolated from gel and sent for Sanger sequencing at Macrogen Europe using primers from Table S1. Editing efficiency was predicted using the TIDE tool ^60^. After confirming successful editing, single cells clones were genotyped as described above.

### CRISPR/Cas9 editing of organoids

PX459 plasmid harboring an sgRNA against mouse Axin1 was purified from E. Coli using the EndoToxin Free Plasmid Maxi Kit (Qiagen). For electroporation, three confluent wells of organoids from a 24-well plate per condition were dissociated with TrypLE express (Gibco). A cell-DNA mixture was prepared by adding 15 µg Cas9 plasmid, 1 µg CMV-GFP plasmid to visualize electroporation efficiency, and up to 100 µl of BTX express buffer (BTX online) containing 10 µM Y-27623 (Sigma). The cell-DNA mixture was transferred to an electroporation cuvette (NEPA) and electroporated using a NEPA21 electroporator (NEPA GENE) with 2× poring pulse (voltage: 175 V, length: 5 ms, interval: 50 ms, polarity: +, decay rate 10) and 5× transfer pulse (voltage: 20 V, length: 50 ms, interval: 50 ms, polarity ±, decay rate 40). Afterwards, cells were incubated for 20 minutes at room temperature in 500 µL opti-MEM (Gibco) containing 10 µM Y-27623, centrifuged at 300 x g at 4 °C and embedded in matrigel. Subsequently, organoids were cultured in EM supplemented with 0.5% Wnt surrogate conditioned medium and 10 μM Y-27623. For selection, 1 μg/μL puromycin (Sigma-Aldrich) was supplemented to the culture medium for 2 days, 2 days after electroporation. To determine the editing efficiency, the organoids were harvested by dissociating the matrigel using 1 mg/mL dispase for 20 minutes at 37 °C. Genotyping was performed as described above using primers in table S1 after which single clones were established by single organoid picking and expansion.

### Antibodies

The following primary antibodies were used for immunoblotting (IB) and immunofluorescence (IF): mouse anti-actin (MP Biomedicals, #691001), mouse anti-GAPDH (Calbiochem, CB1001), mouse anti-β catenin (BD Transduction Laboratories, #610153), rabbit anti-GSK3β (Cell Signaling, #9315), mouse anti-APC (Abcam, ab58), mouse anti-V5 (Genscript, A01724), rabbit anti-V5 (Sigma, V8137), goat anti-AXIN1 (R&D systems, AF3287), rabbit anti-AXIN1 (Cell signaling, #2087), mouse anti-YAP (Santa Cruz, sc-101199). The following secondary antibodies were used for IB and IF: goat-anti-rabbit IrDye680 or IrDye800 (Licor), goat-anti-rabbit Alexa 488 or 568, goat anti-mouse Alexa 488 or 568.

### TOPFlash and 8xGTIIC reporter assays

For the β-catenin-dependent TOPFlash reporter assay, Huh7 were seeded at 100,000 cells per 24-well. After one day, Huh7 cells were transfected with a total amount of 200 ng DNA per well consisting of 30 ng of either TOPFlash or FOPFlash, 20 ng of TK-Renilla, and 150 ng empty vector using FuGene6 (Promega), according to manufacturer’s protocol. HEK293T cells were seeded at 50,000 cells per 24-well. After one day, HEK293T cells were transfected with a total amount of 250 ng DNA per well consisting of 30 ng of either TOPFlash or FOPFlash, 5 ng of TK-Renilla, 50 ng of AXIN1 overexpression construct and 165 ng empty vector using FuGene6. For the TEAD-dependent 8xGTIIC luciferase reporter, cells were seeded at different cell densities and transfected with a total amount of 200 ng DNA per well consisting of 30 ng of 8xGTIIC Luc reporter, 20 ng of thymidine kinase (TK)-Renilla, and 150 ng empty vector using FuGene6. If applicable, 50% Wnt-3a conditioned medium was added 6 hours after transfection. The next day, cells were lysed using 1x Passive lysis buffer (Promega) for 20 minutes at room temperature and 20 μL was transferred to a white costar 96-well plate. Firefly Luciferase and Renilla were measured on a Berthold Luminometer Centro LB960 using the Dual-Luciferase reporter kit (Promega). For siRNA-mediated knock-down, cells were transfected with siRNA to a final concentration of 25nM using lipofectamine RNAiMAX (Thermo) two days prior to transfection with reporter constructs. siRNA AXIN2: Dharmacon L-008809-00-0005. siRNA scrambled: Dharmacon D-001810-01-20.

### Immunofluorescence confocal microscopy

HEK293T cells were seeded in 24-wells onto glass coverslips that were coated with laminin (Sigma-Aldrich) (2,5 μg/mL in PBS) for one hour at 37 °C. The next day, cells were transfected with 100 ng of AXIN1 construct and 150 ng empty vector per well. The following day, cells were fixed in 4% paraformaldehyde (PFA) in 50 mM sodium phosphate buffer, pH 7.4. After 30 minutes, the reaction was quenched by 50mM NH_4_Cl for 30 minutes at room temperature. Next, samples were blocked in blocking buffer containing 2% BSA and 0.1% saponin in PBS for 30 min at room temperature and afterwards incubated with primary antibody in blocking buffer for 1 hour at room temperature. Cells were washed three times with blocking buffer before the addition of secondary antibody and DAPI (Merck) for 45 minutes at room temperature. After three washing steps with blocking buffer, cells were washed with milliQ water and mounted in Prolong Gold (Invitrogen, P36391). Images were acquired with a Zeiss LSM700 confocal microscope and analyzed using ImageJ. Images were quantified using a custom macro for ImageJ, which is provided in Method S1. This macro sets a threshold for signal intensity, subtracts background, and creates cellular outlines based on nuclear localization. Output was manually filtered for incompletely quantified cells.

For detection of β-catenin and YAP in Huh7 and HEK293T, cells were seeded on coverslips and fixed, stained and mounted the following day as described above.

### Immunoprecipitation

For AXIN1 co-immunoprecipitation in HEK293T clones, cells were seeded in five 15 cm dishes (Corning) and cultured until confluency. Cells were resuspended in 5 mL lysis buffer containing 50 mM Tris pH 7.5, 150 mM NaCl, 0.5% Triton X-100, 10% glycerol, 5 mM EDTA, 1 mM DTT, 50 mM sodium, 10 μg/mL aprotinin, 10 μg/mL leupeptin and 100 μM sodium vanadate. Cells were lysed for 1 hour at 4 °C and subsequently centrifuged for 20 minutes at 21130 x g at 4 °C. 45 μL of lysate was collected as total cell lysate. For AXIN1 IP, per sample 1 µg of goat anti-AXIN1 antibody was added, followed by incubation overnight while tumbling at 4 °C. 20 μL of equilibrated protein G-coupled beads (Millipore) was added to each sample, followed by a 2-hour incubation at 4 °C while tumbling. Beads were washed 3 times with lysis buffer, after which proteins were eluted in 2x Sample buffer (SDS, β-mercaptoethanol, glycerol, Tris pH 6.8, bromophenol blue) and boiled for 6 min. For AXIN1 immunoprecipitation in Huh7, JHH5 or JHH6 cells, the same protocol was applied as for HEK293T cells, except that cells were seeded in eight 15 cm dishes (Corning) and resuspended in 500 μL RIPA buffer containing 25 mM Tris pH 7.6, 150 mM NaCl, 1% NP40, 1% deoxycholate, 0,1% SDS supplemented with 1 mM DTT and 50 mM sodium fluoride, aprotinin (10 μg/mL), leupeptin (10 μg/mL) and sodium vanadate (100 μM).

### Western blotting

Proteins were loaded on SDS-PAGE gels and separated on molecular weight. After separation, proteins were transferred to a polyvinylidene fluoride (PVDF) membrane (Immobilon-F) (Millipore). After membrane blocking by Odyssey Blocking Buffer (Li-Cor) diluted 1:1 in Tris-buffered saline (TBS), membranes were incubated with primary antibody overnight at 4 °C in TBS containing 0.05% Tween (TBS-T). After washing with TBS-T, membranes were incubated with secondary antibody for 60 min at RT in TBS-T. After additional washing steps with TBS-T and a final wash with TBS, membranes were scanned by a Typhoon Biomolecular Imager (GE Healthcare).

### Quantitative real-time PCR

Huh7 cells and organoids were lysed in the dish by addition of 350 μL RLT buffer containing 1:100 β-mercaptoethanol. RNA isolation was performed following the RNeasy Kit (Qiagen) protocol, with an additional step to remove genomic DNA by the RNase-free DNase set (Qiagen). Next, cDNA was made from RNA using the iScriptTM cDNA synthesis kit (Biorad). qRT-PCR was performed on a Biorad CFX96 using iQ SYBR Green (Biorad) with primer concentrations of 0.4 nM.

## Acknowledgements

We thank members of the laboratory of MM Maurice for discussions and suggestions. We thank Bart Spee (University Utrecht, Netherlands) for providing Huh7 cells, Ron Smits (Erasmus University, Netherlands) for JHH5 and JHH6 cells and Saskia van Mil (University Medical Center Utrecht, Netherlands) for mouse liver organoids. This work is part of the Oncode Institute, which is partly financed by the Dutch Cancer Society. This work was supported by the ZonMW TOP Grant 91218050 (to MM Maurice), Dutch Cancer Society grant 13112 (to MM Maurice), NWO Gravitation project IMAGINE! (to MM Maurice), and Dutch Cancer Society/TKI-Life Sciences and Health grant 2022-PPS-1/14853 (to MM Maurice).

## Author contributions

AJ Venhuizen: conceptualization, formal analysis, investigation, visualization, methodology, and writing—original draft, review, and editing.

Y van Os: conceptualization, formal analysis, investigation, visualization and methodology.

ML Kaptein: conceptualization, formal analysis, investigation, visualization and methodology.

MT Aarts: conceptualization, formal analysis, investigation, visualization and methodology.

D Xanthakis: formal analysis, investigation and methodology

I Jordens: conceptualization, formal analysis, investigation, visualization, methodology, and writing—original draft, review, and editing.

MM Maurice: formal analysis, supervision, funding acquisition, investigation, visualization, methodology, and writing—original draft, review, and editing.

## Supplemental information

Document S1: Figures S1–S8, Table S1, Method S1

