## Supplemental document S1 for "Hepatocellular carcinoma-associated *AXIN1* mutations drive low levels of Wnt/β-catenin pathway activity that allow for niche-independent growth and YAP/TAZ signaling"

**Document S1 – Supplemental information**

Figure S1

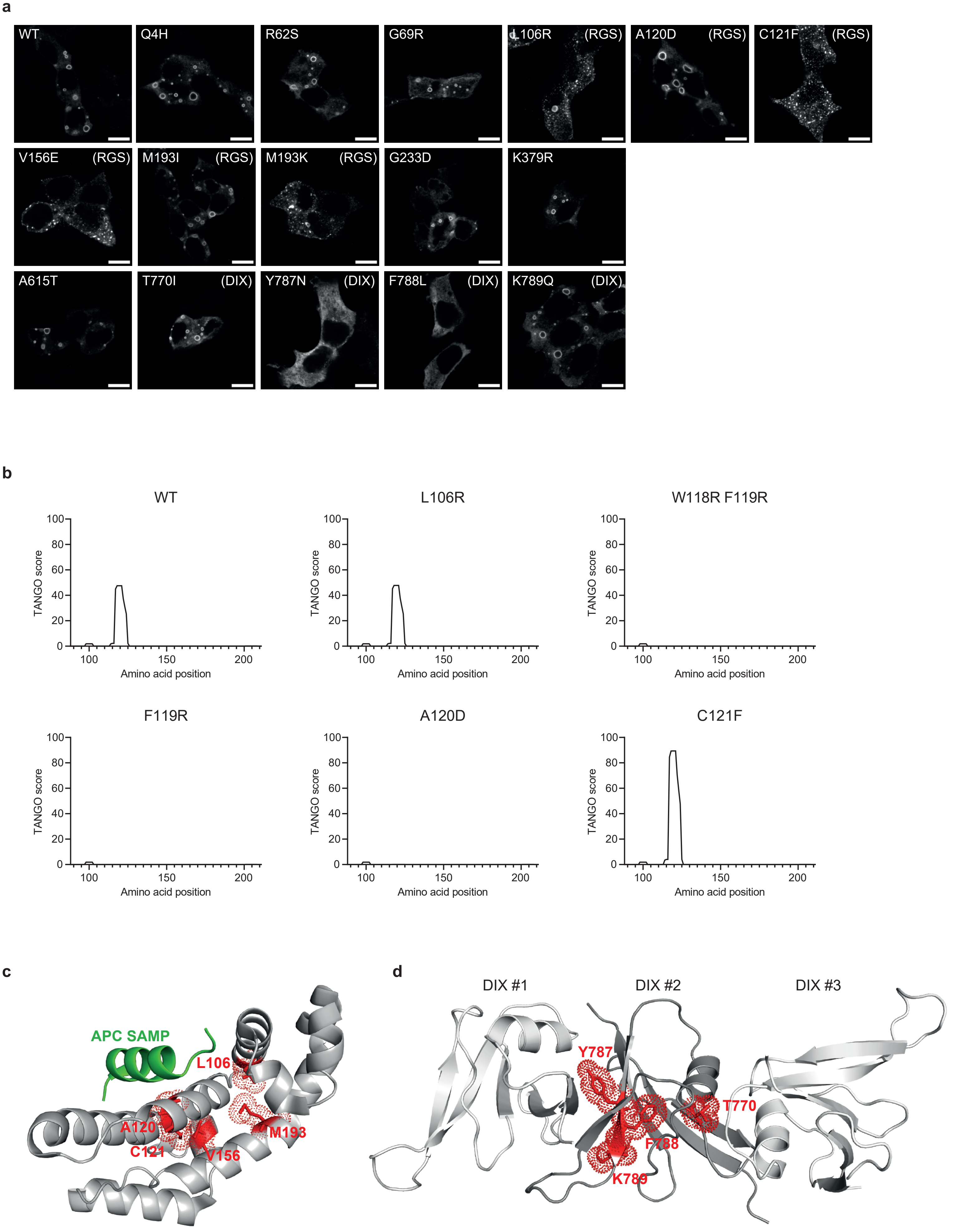
a
WT
Q4H
R62S
G69R
L106R (RGS)
A120D (RGS)
C121F (RGS)
V156E (RGS)
M193I (RGS)
M193K (RGS)
G233D
K379R
A615T
T770I (DIX)
Y787N (DIX)
F788L (DIX)
K789Q (DIX)
b
WT
L106R
W118R F119R
F119R
A120D
C121F
TANGO score
Amino acid position
0
20
40
60
80
100
150
200
c
APC SAMP
L106
A120
C121
V156
M193
d
DIX #1
DIX #2
DIX #3
Y787
T770
F788
K789

### Figure S2

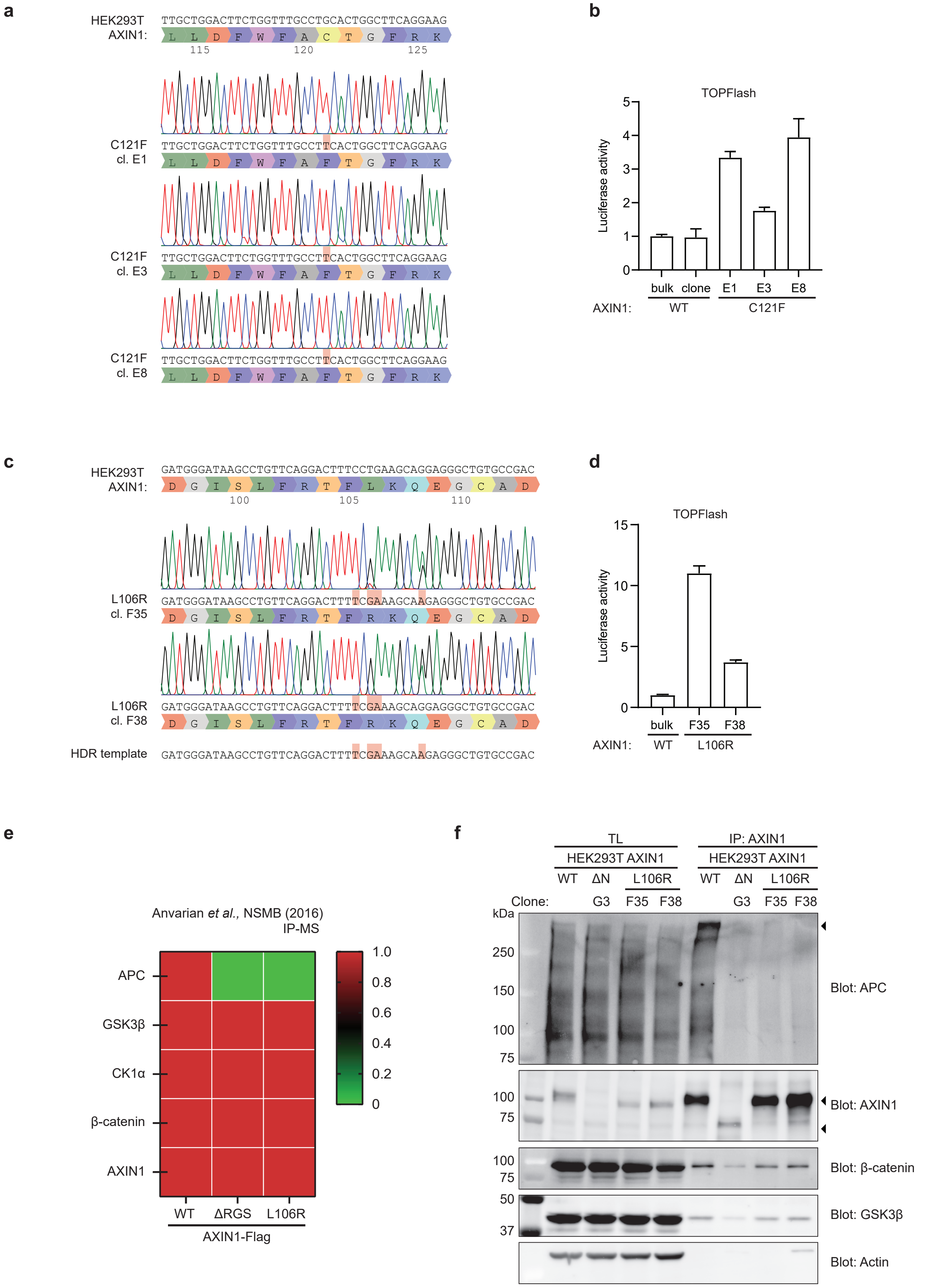

a
HEK293T AXIN1:
TTGCTGGACTTCTGGTTTGCCTGCACTGGCTTCAGGAAG
L L D F W F A C T G F R K
115 120 125
C121F cl. E1
TTGCTGGACTTCTGGTTTGCCTTCACTGGCTTCAGGAAG
L L D F W F A F T G F R K
C121F cl. E3
TTGCTGGACTTCTGGTTTGCCTTCACTGGCTTCAGGAAG
L L D F W F A F T G F R K
C121F cl. E8
TTGCTGGACTTCTGGTTTGCCTTCACTGGCTTCAGGAAG
L L D F W F A F T G F R K
b
TOPFlash
Luciferase activity
bulk clone E1 E3 E8
AXIN1: WT C121F
c
HEK293T AXIN1:
GATGGGATAAGCCTGTTCAGGACTTTCCTGAAGCAGGAGGGCTGTGCCGAC
D G I S L F R T F L K Q E G C A D
100 105 110
L106R cl. F35
GATGGGATAAGCCTGTTCAGGACTTTTCGAAAGCAAGAGGGCTGTGCCGAC
D G I S L F R T F R K Q E G C A D
L106R cl. F38
GATGGGATAAGCCTGTTCAGGACTTTTCGAAAGCAGGAGGGCTGTGCCGAC
D G I S L F R T F R K Q E G C A D
HDR template
GATGGGATAAGCCTGTTCAGGACTTTTCGAAAGCAAGAGGGCTGTGCCGAC
d
TOPFlash
Luciferase activity
bulk F35 F38
AXIN1: WT L106R
e
Anvarian et al., NSMB (2016) IP-MS
APC
GSK3β
CK1α
β-catenin
AXIN1
WT ΔRGS L106R
AXIN1-Flag
1.0 0.8 0.6 0.4 0.2 0
f
TL
HEK293T AXIN1
WT ΔN L106R
IP: AXIN1
HEK293T AXIN1
WT ΔN L106R
Clone: G3 F35 F38 G3 F35 F38
kDa
250 150 100 75
Blot: APC
100 75
Blot: AXIN1
100 75
Blot: β-catenin
50 37
Blot: GSK3β
Blot: Actin

Figure S3

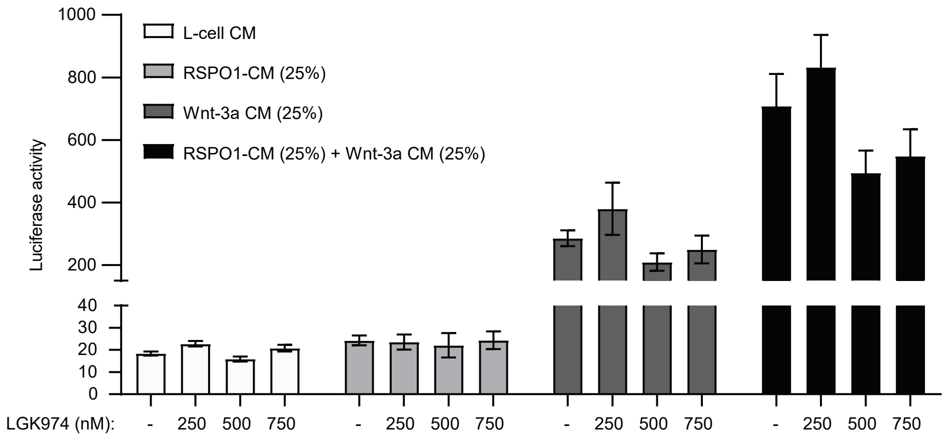

L-cell CM
RSPO1-CM (25%)
Wnt-3a CM (25%)
RSPO1-CM (25%) + Wnt-3a CM (25%)
Luciferase activity
1000
800
600
400
200
40
30
20
10
0
LGK974 (nM):
- 250 500 750
- 250 500 750
- 250 500 750
- 250 500 750

Figure S4

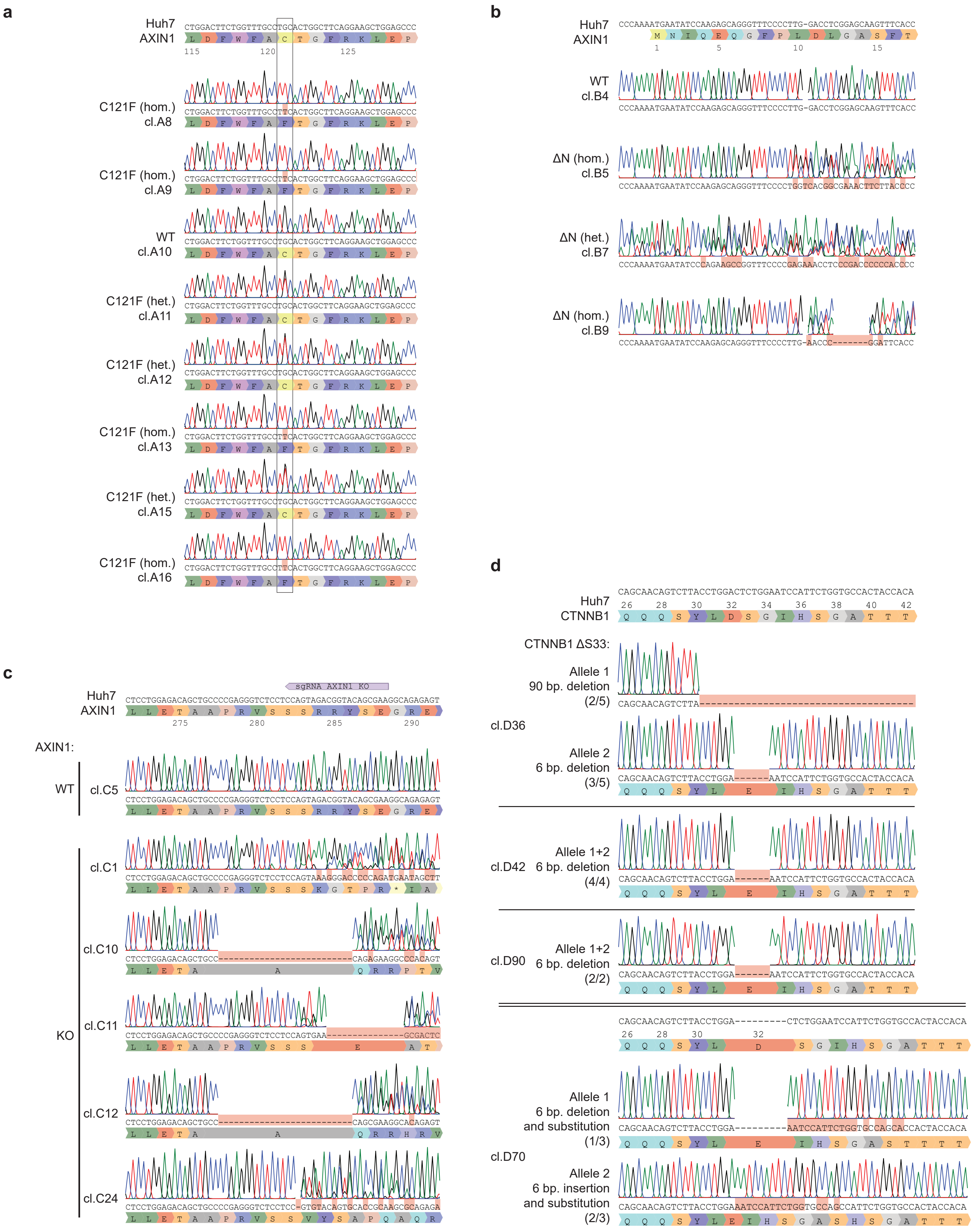

a
Huh7
AXIN1
C121F (hom.)
cl.A8
C121F (hom.)
cl.A9
WT
cl.A10
C121F (het.)
cl.A11
C121F (het.)
cl.A12
C121F (hom.)
cl.A13
C121F (het.)
cl.A15
C121F (hom.)
cl.A16
b
Huh7
AXIN1
WT
cl.B4
ΔN (hom.)
cl.B5
ΔN (het.)
cl.B7
ΔN (hom.)
cl.B9
c
sgRNA AXIN1 KO
Huh7
AXIN1
AXIN1:
WT
cl.C5
KO
cl.C1
cl.C10
cl.C11
cl.C12
cl.C24
d
Huh7
CTNNB1
CTNNB1 ΔS33:
Allele 1
90 bp. deletion
(2/5)
cl.D36
Allele 2
6 bp. deletion
(3/5)
Allele 1+2
cl.D42 6 bp. deletion
(4/4)
Allele 1+2
cl.D90 6 bp. deletion
(2/2)
Allele 1
6 bp. deletion
and substitution
(1/3)
cl.D70
Allele 2
6 bp. insertion
and substitution
(2/3)

### Figure S5

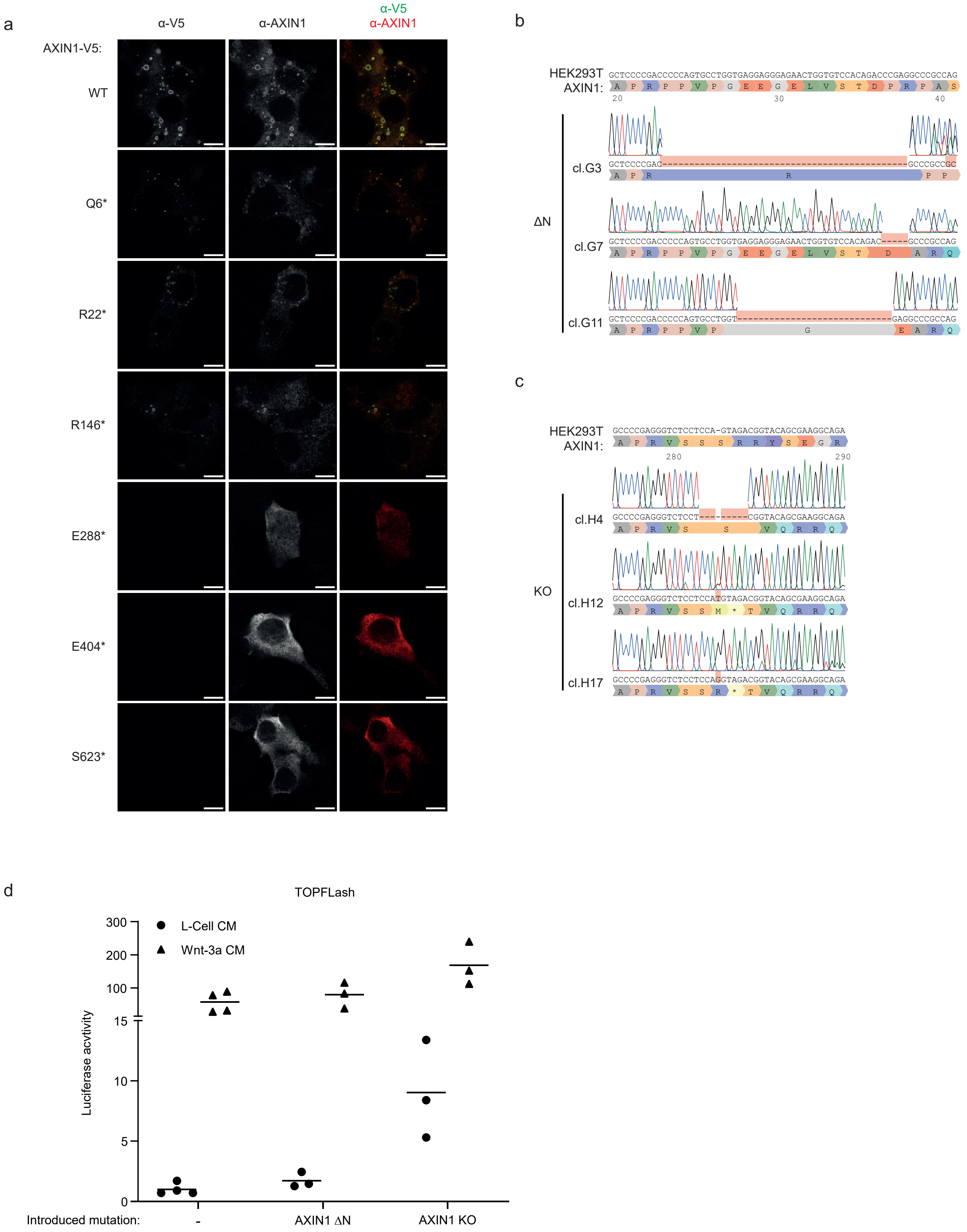

a
α-V5
α-AXIN1
α-V5
α-AXIN1
AXIN1-V5:
WT
Q6*
R22*
R146*
E288*
E404*
S623*
b
HEK293T
AXIN1:
ΔN
cl.G3
cl.G7
cl.G11
c
HEK293T
AXIN1:
KO
cl.H4
cl.H12
cl.H17
d
TOPFLash
L-Cell CM
Wnt-3a CM
Luciferase acvtivity
300
200
100
15
10
5
0
Introduced mutation:
-
AXIN1 ΔN
AXIN1 KO

Figure S6

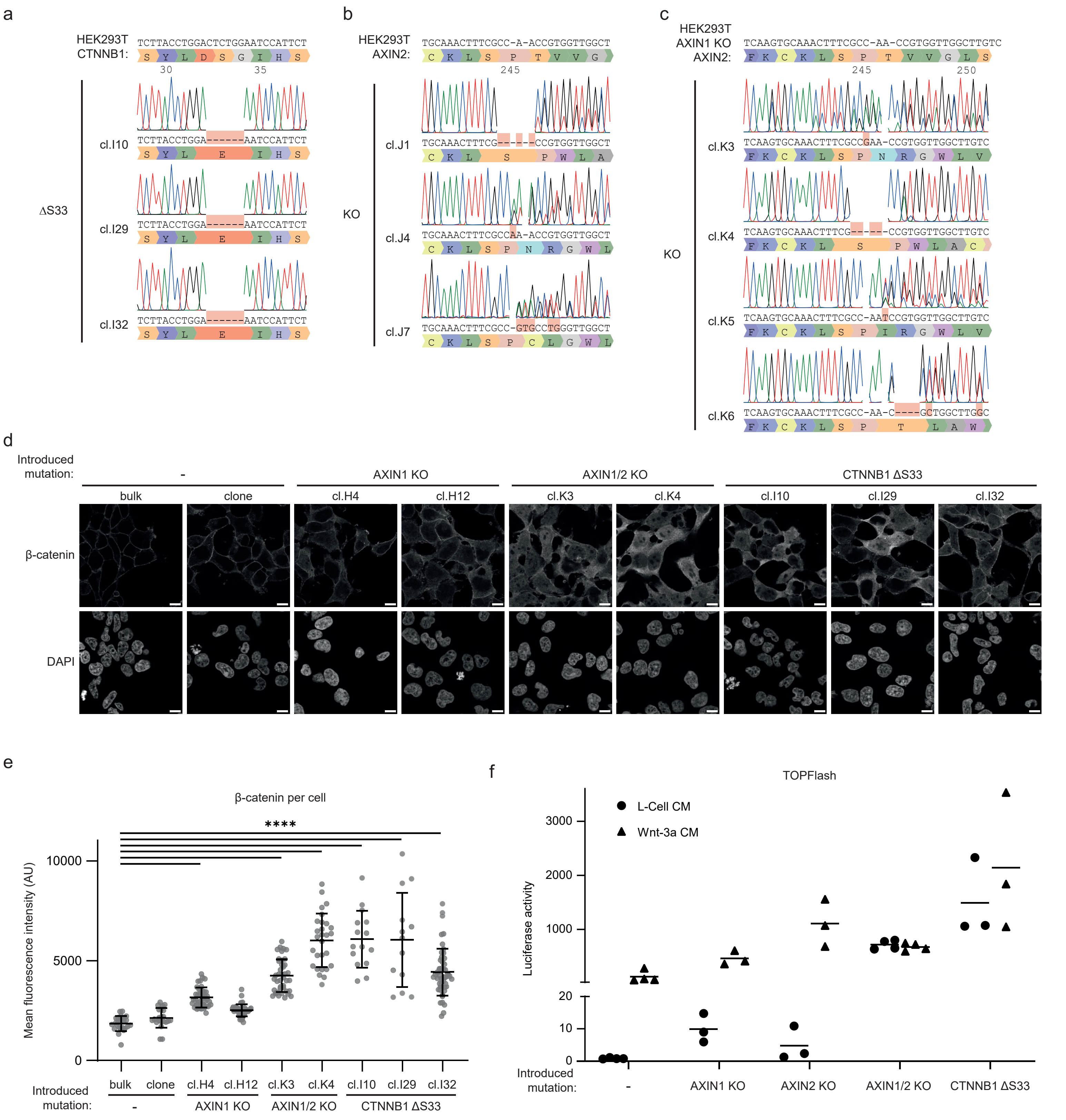

a
HEK293T CTNNB1:
TCTTACCTGGACTCTGGAATCCATTCT
ΔS33
cl.I10
cl.I29
cl.I32
TCTTACCTGGA------AATCCATTCT
b
HEK293T AXIN2:
TGCAAACTTTCGCC-A-ACCGTGGTTGGCT
KO
cl.J1
cl.J4
cl.J7
c
HEK293T AXIN1 KO AXIN2:
TCAAGTGCAAACTTTCGCC-AA-CCGTGGTTGGCTTGTC
KO
cl.K3
cl.K4
cl.K5
cl.K6
d
Introduced mutation:
-
AXIN1 KO
AXIN1/2 KO
CTNNB1 ΔS33
bulk
clone
cl.H4
cl.H12
cl.K3
cl.K4
cl.I10
cl.I29
cl.I32
β-catenin
DAPI
e
β-catenin per cell
****
Mean fluorescence intensity (AU)
Introduced mutation:
f
TOPFlash
L-Cell CM
Wnt-3a CM
Luciferase activity
AXIN2 KO

Figure S7

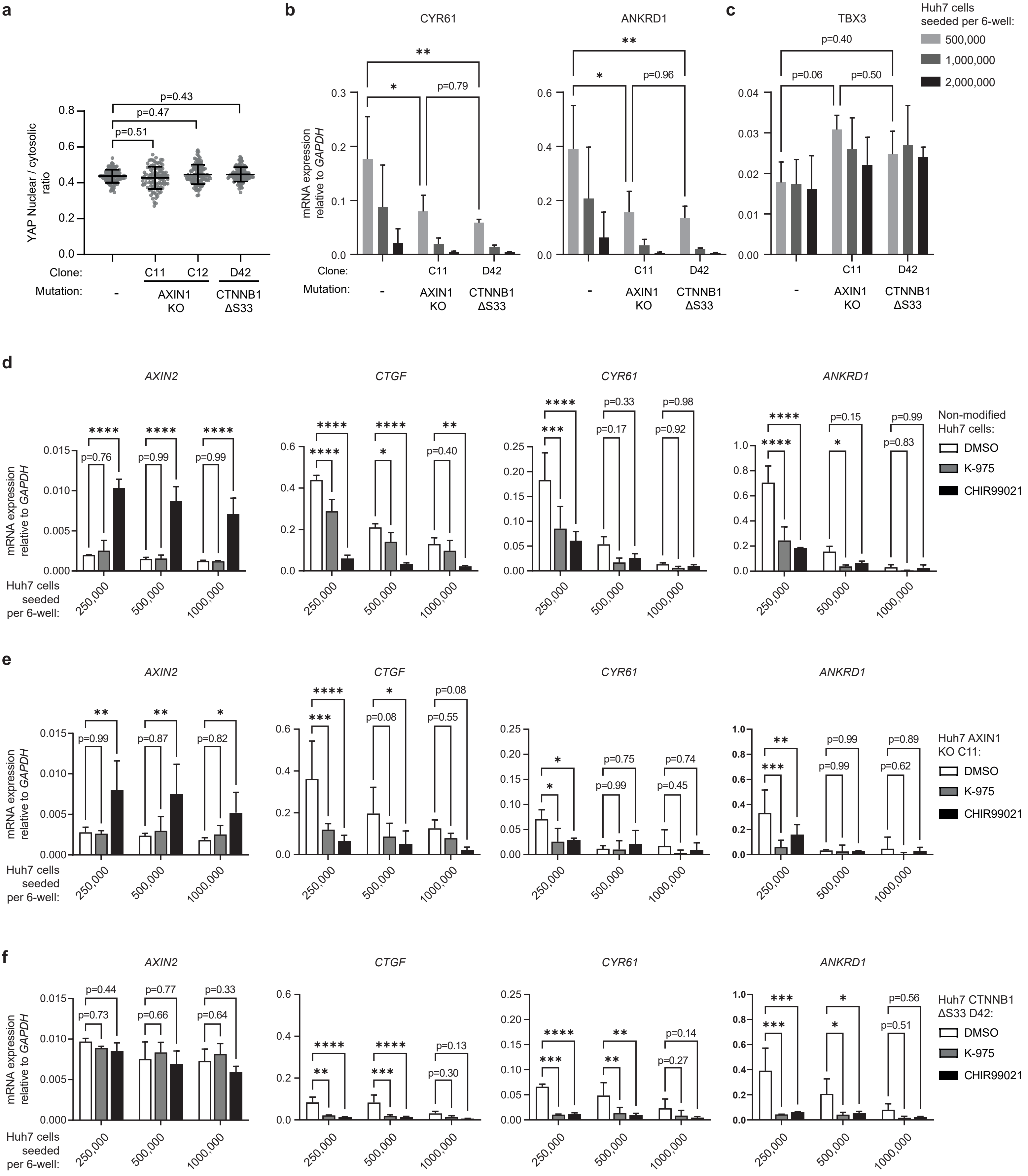

a
YAP Nuclear / cytosolic ratio
p=0.43
p=0.47
p=0.51
Clone: C11 C12 D42
Mutation: - AXIN1 KO CTNNB1 ΔS33
b
CYR61
ANKRD1
mRNA expression relative to GAPDH
**
*
p=0.79
p=0.96
Clone: C11 D42
Mutation: - AXIN1 KO CTNNB1 ΔS33
c
TBX3
p=0.40
p=0.06
p=0.50
Huh7 cells seeded per 6-well:
500,000
1,000,000
2,000,000
d
AXIN2
CTGF
CYR61
ANKRD1
Non-modified Huh7 cells:
DMSO
K-975
CHIR99021
Huh7 cells seeded per 6-well:
250,000
500,000
1000,000
e
Huh7 AXIN1 KO C11:
f
Huh7 CTNNB1 ΔS33 D42:

### Figure S8

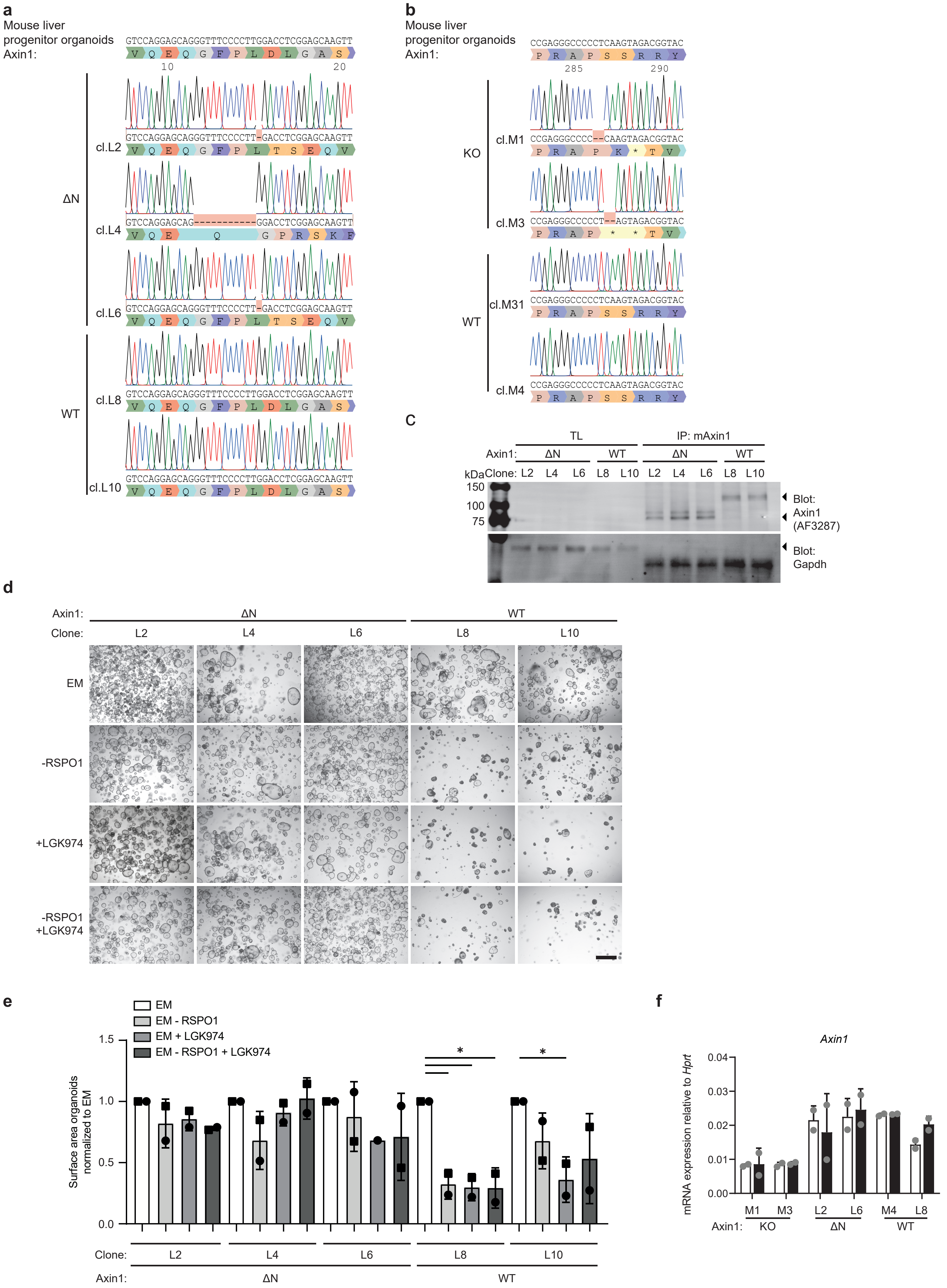
a
Mouse liver progenitor organoids Axin1:
GTCCAGGAGCAGGGTTTCCCCTTGGACCTCGGAGCAAGTT
ΔN
cl.L2
GTCCAGGAGCAGGGTTTCCCCTT-GACCTCGGAGCAAGTT
cl.L4
GTCCAGGAGCAG----------GGACCTCGGAGCAAGTT
cl.L6
GTCCAGGAGCAGGGTTTCCCCTT-GACCTCGGAGCAAGTT
WT
cl.L8
GTCCAGGAGCAGGGTTTCCCCTTGGACCTCGGAGCAAGTT
cl.L10
GTCCAGGAGCAGGGTTTCCCCTTGGACCTCGGAGCAAGTT
b
Mouse liver progenitor organoids Axin1:
CCGAGGGCCCCCTCAAGTAGACGGTAC
KO
cl.M1
CCGAGGGCCCC--CAAGTAGACGGTAC
cl.M3
CCGAGGGCCCCCT--AGTAGACGGTAC
WT
cl.M31
CCGAGGGCCCCCTCAAGTAGACGGTAC
cl.M4
CCGAGGGCCCCCTCAAGTAGACGGTAC
c
TL
IP: mAxin1
Axin1: ΔN WT ΔN WT
kDa Clone: L2 L4 L6 L8 L10 L2 L4 L6 L8 L10
150
100
75
Blot: Axin1 (AF3287)
Blot: Gapdh
d
Axin1: ΔN WT
Clone: L2 L4 L6 L8 L10
EM
-RSPO1
+LGK974
-RSPO1 +LGK974
e
EM
EM - RSPO1
EM + LGK974
EM - RSPO1 + LGK974
Surface area organoids normalized to EM
1.5
1.0
0.5
0.0
Clone: L2 L4 L6 L8 L10
Axin1: ΔN WT
f
Axin1
mRNA expression relative to Hprt
0.04
0.03
0.02
0.01
0.00
M1 M3 L2 L6 M4 L8
Axin1: KO ΔN WT

**Supplemental figure S1. Related to figure 1 - Missense mutations within the RGS domain of AXIN1 drive Wnt signaling. (a)** Representative immunofluorescence images of HEK293T cells overexpressing the indicated AXIN1-V5 variants. Fixed cells were labeled for V5. Scale bar represents 15 µm. **(b)** Aggregation propensity of indicated AXIN1 RGS variants predicted by TANGO. Cancer mutation A120D is predicted to function as rescue mutation for RGS aggregation, correlating with the observed puncta formation of this variant in (a). Cancer mutation C121F increases the aggregation propensity of the RGS domain, explaining why this cancer mutation shows stronger Wnt pathway activation than the L106R mutation. **(c)** Superimposition of residues mutated (red) in HCC onto the crystal structure of rat AXIN1 RGS (pdb: 1EMU). AXIN1-binding SAMP domain of APC is depicted in green. **(d)** Superimposition of residues mutated (red) in HCC onto the crystal structure of 3 polymerized rat AXIN1 DIX domains (pdb: 1WSP).

**Supplemental figure S2. Related to figure 1 - Missense mutations within the RGS domain of AXIN1 drive Wnt/β-catenin signaling in HEK293T cells. (a)** Sanger sequencing results of HEK293T clones harboring the AXIN1 C121F mutation introduced by prime editing. **(b)** β-catenin-dependent TOPFlash assay of HEK293T clones harboring the C121F mutation. The non-modified clone underwent the same procedure, but did not acquire the mutation. Graph shows a representative experiment (N=3) with mean +/- SD of n=2 wells. **(c)** Sanger sequencing results of HEK293T clones harboring the AXIN1 L106R mutation introduced via homologous recombination. **(d)** TOPFlash assay of HEK293T clones harboring the L106R mutation. Graph shows a representative experiment (N=3) with mean +/- SD of n=2 wells. **(e)** Heatmap displaying the relative amounts of β-catenin destruction complex members bound by FLAG-tagged AXIN1 mutants, as previously published[15]. **(f)** Co-immunoprecipitation experiment of endogenous AXIN1 (using AF3287 antibody) in indicated *AXIN1*-mutant HEK293T cell lines. Actin is used as loading control.

**Supplemental figure S3. Related to figure 1 – Wnt/β-catenin pathway activity in Huh7 cells.** TOPFlash reporter assay of Huh7 cells treated with Wnt-3a conditioned medium, RSPO1 conditioned medium, or L-cell conditioned medium as control. In addition, cells were treated with increasing concentrations of porcupine inhibitor LGK974. Graph shows a representative experiment (N=3) with mean +/- SD of n=2 wells.

**Supplemental figure S4. Related to figure 1, 2, 3 and 4 - Sanger sequencing of genetically modified Huh7 cells.** Sanger sequencing results of Huh7 **(a)** AXIN1 C121F, **(b)** AXIN1 ΔN, **(c)**

AXIN1 KO and **(d)** β-catenin ΔS33. For the β-catenin ΔS33 clones in (d), the frequency of each allele was determined by subcloning in pJET. Numbers between brackets indicate the amount of sanger sequencing reads found for each type of mutation.

**Supplemental figure S5. Related to figure 2 - Frameshift mutations in 5' coding regions yield an N-terminally truncated AXIN1 variant with partially retained functionality. (a)** Representative immunofluorescence images of HEK293T cells overexpressing different AXIN1 truncating variants. Fixed cells were stained for V5 and AXIN1. **(b)** Sanger sequencing results of HEK293T cells harboring AXIN1 ΔN and **(c)** AXIN1 KO. **(d)** TOPFlash reporter assay comparing non-modified HEK293T cells with clones harboring AXIN1 ΔN and KO mutations. Graph shows a representative experiment (N=3), where one dot represents the mean of technical duplicates of one clone. Cells were treated with Wnt-3a conditioned medium or L-cell conditioned medium as control.

**Supplemental figure S6. Related to figure 3 - *AXIN1*-mutant cells have moderate Wnt/β-catenin signaling levels, while *CTNNB1*-mutant cells are Wnt-high.** Sanger sequencing results of HEK293T **(a)** β-catenin ΔS33, **(b)** AXIN2 KO, **(c)** as well as AXIN1 and AXIN2 KO. **(d)** Representative immunofluorescence images of HEK293T cells harboring different Wnt pathway mutations, labeled for β-catenin and DAPI. Scale bar represents 15 µm. **(e)** Quantification of immunofluorescence images from (d). Quantified using an automated ImageJ script as described in the methods section. Significance was determined using one-way ANOVA. **** indicates $p \leq 0.0001$. **(f)** TOPFlash reporter assay comparing non-modified HEK293T cells to clones harboring different Wnt pathway mutations. Graph shows a representative experiment (N=3), where one dot represents the mean of technical duplicates of one clone. Cells were treated with Wnt-3a conditioned medium or L-cell conditioned medium as control.

**Supplemental figure S7. Related to figure 4 - Wnt pathway activation leads to dose-dependent inhibition of YAP/TAZ signaling in Huh7 cells. (a)** Quantification of immunofluorescence images belonging to experiment in Fig. 4d. Quantified using an automated ImageJ script. Per cell, nuclear intensity levels were divided by the total cytosolic intensity to acquire a nuclear/cytosolic YAP ratio. One-way ANOVA was performed to determine significance. * indicates $p \leq 0.05$, **** indicates $p \leq 0.0001$. Non-significant comparisons were left out for clarity. **(b-c)** RT-qPCR depicting expression of YAP/TAZ target genes (b) *CYR61* and *ANKRD1* and (c) *TBX3* relative to the household gene *GAPDH* for different Huh7 clones at increasing cell density. Two-way ANOVA was performed to determine significance. **(d-f)** Quantitative polymerase chain reaction experiments for (d) non-modified, (e) AXIN1-deficient or (f) β-catenin-mutant Huh7 depicting expression of *AXIN2, CTGF* and *CYR61 and ANKRD1* relative to the household gene *GAPDH* for Huh7 cells treated with different small molecules at increasing cell density. Bars and error bars represent the mean and standard deviation of three biological replicates. Two-way ANOVA was performed to determine significance. * indicates p ≤ 0.05, ** indicates p ≤ 0.01, *** indicates p ≤ 0.001, **** indicates p ≤ 0.0001.

**Supplemental figure S8. Related to figure 5 - Axin1 mutations promote Wnt ligand-independent growth in mouse liver progenitor organoids. (a)** Sanger sequencing results of mouse liver progenitor organoids harboring frameshifts leading to Axin1 ΔN clones and **(b)** Axin1 KO. **(c)** Western blot of immunoprecipitated Axin1 from WT and Axin1 ΔN mouse liver progenitor organoids. Arrows indicate the WT and truncated Axin1. Gapdh was used as loading control, indicated by arrow. **(d)** Representative brightfield images of liver progenitor organoids cultured in expansion medium (EM) with and without 10% RSPO1 conditioned medium and 500 nM LGK974. See figure 5b for the diagram indicating the protocol used. The scale bar represents 500 µm. **(e)** Quantification of biological replicates as performed in (d). Quantification was performed by determining cell surface area of the organoids using OrganoSeg analysis software[61]. Each condition was normalized to organoids cultured in EM. Significance was determined using one-way ANOVA. * indicates p ≤ 0.05. EM, Expansion medium. **(f)** RT-qPCR experiments for *Axin1* mRNA levels relative to *Hprt* after treatment with LGK974 or DMSO as control, similar to figures 5g-j. Bars and error bars represent the mean of two biological replicates +/- SD. Given only N=2, the values for the biological replicates are depicted in grey.

| **Table S1** | |
|---|---|
| **CRISPR/Cas9 oligos (including overhang)** | |
| sgRNA hAXIN1 5' exon 2 fw | caccgAACTTGCTCCGAGGTCCAAG |
| sgRNA hAXIN1 5' exon 2 rv | aaacCTTGGACCTCGGAGCAAGTT |
| sgRNA hAXIN1 3' exon 2 fw | caccgTTCGCTGTACCGTCTACTGG |
| sgRNA hAXIN1 3' exon 2 rv | aaacCCAGTAGACGGTACAGCGAAC |
| sgRNA hAXIN1 L106R fw | caccGGTTCAGGACTTTCCTGAAGC |
| sgRNA hAXIN1 L106R rv | aaacGCTTCAGGAAAGTCCTGAACC |
| ssDNA for hAXIN1 L106R knock-in (* denotes phosphorothioate bonds) | *T*CTTCTCCTCGTTCGAGTCACAGGGCTCCAGCTTCCTGAAGCCAGTGCAGGCAAACCAGAAGTCCAGCAAGTCGGCACAGCCCTCTTGCTTTCGAAAAGTCCTGAACAGGCTTATCCCATCTT*G*G |
| sgRNA hAXIN2 3' exon 2 fw | caccGTGCAAACTTTCGCCAACCG |
| sgRNA hAXIN2 3' exon 2 rv | aaacCGGTTGGCGAAAGTTTGCAC |
| sgRNA mAXIN1 5' exon 2 fw | caccGCTTTCCCAGAACAGAAACTG |
| sgRNA mAXIN1 5' exon 2 rv | aaacCAGTTTCTGTTCTGGGAAAGC |
| sgRNA mAXIN1 3' exon 2 fw | caccGTTGTACCGTCTACTTGAGG |
| sgRNA mAXIN1 3' exon 2 rv | aaacCCTCAAGTAGACGGTACAAC |
| **Prime editing oligos (including overhang)** | |
| hCTNNB ΔS33 pegRNA spacer (excluding overhangs) | CAACAGTCTTACCTGGACTC |
| hCTNNB ΔS33 3’ extension sequence (excluding overhangs) | TGGCACCAGAATGGATTTCCAGGTAAGAC |
| hCTNNB ΔS33 Nicking sgRNA (PE3) (excluding overhangs) | CCACTCATACAGGACTTGGG |
| hAXIN1 C121F pegRNA spacer fw | caccGCAGCTTCCTGAAGCCAGTGCgtttt |
| hAXIN1 C121F pegRNA spacer rv | ctctaaaacGCACTGGCTTCAGGAAGCTGC |
| pegRNA scaffold fw | AGAGCTAGAAATAGCAAGTTAAAATAAGGCTAGTCCGTTATCAACTTGAAAAAGTGGCACCGAGTCG |
| pegRNA scaffold rv | GCACCGACTCGGTGCCACTTTTTCAAGTTGATAACGGACTAGCCTTATTTTAACTTGCTATTTCTAG |
| hAXIN1 C121F pegRNA extension fw | gtgcTGGTTTGCCTtCACTGGCTTCAGGAAG |
| hAXIN1 C121F pegRNA extension rv | aaaaCTTCCTGAAGCCAGTGaAGGCAAACCA |
| hAXIN1-C121F Nicking sgRNA (PE3b) fw | caccGACTTCTGGTTTGCCTtCAC |
| hAXIN1-C121F Nicking sgRNA (PE3b) rv | aaacGTGaAGGCAAACCAGAAGTC |
| **Genotyping primers** | |
| gPCR hAXIN1 5' exon 2 fw | GCGTCATCGTGAGTCTTGTC |
| gPCR hAXIN1 5' exon 2 rv | TGTCTCCAGGAGCAGCTT |
| gPCR hAXIN1 5' exon 2 fw | CCCCCACCCCACCATACTTGAA |
| gPCR hAXIN1 5' exon 2 rv | TCATCAGCACCTTTCCCTGGCT |

| | |
|---|---|
| gPCR hAXIN2 5' exon 2 fw | GCTGAAGCCTGCCACCAAGACC |
| gPCR hAXIN2 5' exon 2 rv | CCACAACCCAGCTGCCTCCCTA |
| hCTNNB1 ΔS33 fw | GCCTGGATGCAGTACCATTCTTCCAC |
| hCTNNB1 ΔS33 rv | AGACACCATCTGAGGAGAACGCA |
| hCTNNB1 ΔS33 sequencing primer | ACACTCACTATCCACAGTTCAGCA |
| gPCR mAxin1 5' exon 2 fw | AGATGTCCTCCATGACTCAGGCT |
| gPCR mAxin1 5' exon 2 rv | ACCCTGAGCTCTGGTCACTGCA |
| gPCR mAxin1 3' exon 2 fw | AGCCACCCCAAGACGTTCAGAT |
| gPCR mAxin1 3' exon 2 rv | TCCTTGCTCCTTTGCCAGGTCT |
| **qRT-PCR primers** | |
| hGAPDH fw | CTTTTGCGTCGCCAG |
| hGAPDH rv | TTGATGGCAACAATATCCAC |
| hAXIN1 fw | CCTGTGGTCTACCCGTGTCT |
| hAXIN1 rv | GCTATGAGGAGTGGTCCAGG |
| hAXIN2 fw | AAAGAGAGGAGGTTCAGATG |
| hAXIN2 rv | CTGAGTCTGGGAATTTTTCTTC |
| hLGR5 fw | GGTGACAACAGCAGTATGGACGA |
| hLGR5 rv | GAAGGTGAACACTGCACTGAATGAA |
| hCYR61 fw | GATCTGCAGAGCTCAGTCAGAG |
| hCYR61 rv | CCATCAATACATGTGCACTG |
| hCTGF fw | CTGGAAGAGAACATTAAGAAGG |
| hCTGF rv | GGTATGTCTTCATGCTGGTG |
| hAREG fw | GAGCCGACTATGACTACTCAGA |
| hAREG rv | TCACTTTCCGTCTTGTTTTGGG |
| hANRKD1 fw | AGTAGAGGAACTGGTCACTGG |
| hANRKD1 rv | TGTTTCTCGCTTTTCCACTGTT |
| hTBX3_fw | AGCGATCACGCAACGTGGCA |
| hTBX3_rv | GGCTTCGCTGGGACACAGATCTTT |
| mHprt_fw | AAGCTTGCTGGTGAAAAGGA |
| mHprt_rv | TTGCGCTCATCTTAGGCTTT |
| mAxin1_fw | ACCCAGTACCACAGAGGACG |
| mAxin1_rv | CTGCTTCCTCAACCCAGAAG |
| mAxin2_fw | GGACTGGGGAGCCTAAAGGT |
| mAxin2_rv | AAGGAGGGACTCCATCTACGC |
| mLgr5_fw | AGAACACTGACTTTGAATGG |
| mLgr5_rv | CACTTGGAGATTAGGTAACTG |
| mAlb_fw | GCGCAGATGACAGGGCGGAA |
| mAlb_rv | GTGCCGTAGCATGCGGGAGG |

### Method S1 – Macro ImageJ analysis

```
DAPI = "1"
bCat = "2"

title = getTitle();
run("Duplicate...", "duplicate channels=" + DAPI);
DAPIdup = getTitle();
run("Gaussian Blur...", "sigma=10");
setAutoThreshold("Default dark");
setOption("BlackBackground", false);
run("Convert to Mask");

selectImage(title);
run("Select None");
run("Duplicate...", "duplicate channels=" + bCat);
bCatdup = getTitle();
run("Median...", "radius=20");
setAutoThreshold("Minimum dark");
waitForUser("ok")
run("Convert to Mask");
selectImage(title);
run("Select None");
run("Duplicate...", "duplicate channels=" + DAPI);
run("Gaussian Blur...", "sigma=20");
run("Find Maxima...", "prominence=20 output=[Segmented Particles]");
segmentdup = getTitle();
imageCalculator("Min create", bCatdup,segmentdup);
outlinedup = getTitle();
run("Analyze Particles...", "clear add");
imageCalculator("Min create", bCatdup,segmentdup);
selectImage(DAPIdup);
run("Select None");
run("Invert");
imageCalculator("Min create", outlinedup, DAPIdup);
cytodup = getTitle();

roiManager("Show All");
waitForUser("please check valid cytosols");
roiManager("Show None");
run("Select None");
nROIs = roiManager("count");
for (i = 0; i < nROIs; i++) {
		selectImage(cytodup);
		run("Duplicate...", " ");
		run("Invert");
		roiManager("Select", i);
		run("Clear Outside");
		run("Select None");
		run("Invert");
		run("Convert to Mask");
		run("Create Selection");
		selectImage(title);
		Stack.setChannel(bCat);
		run("Restore Selection");
		//waitForUser("OK");
		run("Measure");
		bottomrow = (nResults-1);
		print(bottomrow);
		area = getResult("Area", bottomrow);
		print(area);
		//waitForUser("OK");
		if (area<100){
				IJ.deleteRows(bottomrow, bottomrow);
		} else if (area>1000){
				IJ.deleteRows(bottomrow, bottomrow);
		}
		//waitForUser("OK");
		run("Select None");
		selectImage(DAPIdup);
		run("Select None");
		run("Duplicate...", " ");
		roiManager("Select", i);
		run("Clear Outside");
		run("Select None");
		run("Invert");
		run("Convert to Mask");
		run("Create Selection");
		//waitForUser("OK");
		selectImage(title);
		Stack.setChannel(bCat);
		run("Restore Selection");
		run("Measure");
		bottomrow = (nResults-1);
		if (area<100){
				IJ.deleteRows(bottomrow, bottomrow);
		} else if (area>1000){
				IJ.deleteRows(bottomrow, bottomrow);
		}
		//waitForUser("OK");
}
waitForUser("please check cytosoldata");
```
